# Oculomotor plant and neural dynamics suggest gaze control requires integration on distributed timescales

**DOI:** 10.1101/2021.06.30.450653

**Authors:** Andrew Miri, Brandon J. Bhasin, Emre R. F. Aksay, David W. Tank, Mark S. Goldman

## Abstract

A fundamental principle of biological motor control is that the neural commands driving movement must conform to the response properties of the motor plants they control. In the oculomotor system, characterizations of oculomotor plant dynamics traditionally supported models in which the plant responds to neural drive to extraocular muscles on exclusively short, subsecond timescales. These models predict that the stabilization of gaze during fixations between saccades requires neural drive that approximates eye position on longer timescales and is generated through the temporal integration of brief eye velocity-encoding signals that cause saccades. However, recent measurements of oculomotor plant behaviour have revealed responses on longer timescales. Furthermore, measurements of firing patterns in the oculomotor integrator have revealed a more complex encoding of eye movement dynamics. Yet, the link between these observations has remained unclear. Here we use measurements from the larval zebrafish to link dynamics in the oculomotor plant to dynamics in the neural integrator. The oculomotor plant in both anaesthetized and awake larval zebrafish was characterized by a broad distribution of response timescales, including those much longer than one second. Analysis of the firing patterns of oculomotor integrator neurons, which exhibited a broadly distributed range of decay time constants, demonstrates the sufficiency of this activity for stabilizing gaze given an oculomotor plant with distributed response timescales. This work suggests that leaky integration on multiple, distributed timescales by the oculomotor integrator reflects an inverse model for generating oculomotor commands, and that multi-timescale dynamics may be a general feature of motor circuitry.

**KEY POINTS:**

- Recent observations of oculomotor plant response properties and neural activity across the oculomotor system have called into question classical formulations of both the oculomotor plant and the oculomotor integrator.
- Here we use measurements from new and published experiments in the larval zebrafish together with modelling to reconcile recent oculomotor plant observations with oculomotor integrator function.
- We developed computational techniques to characterize oculomotor plant responses over several seconds in awake animals, demonstrating that long timescale responses seen in anesthetized animals extend to the awake state.
- Analysis of firing patterns of oculomotor integrator neurons demonstrates the sufficiency of this activity for stabilizing gaze given an oculomotor plant with multiple, distributed response timescales.
- Our results support a formulation of gaze stabilization by the oculomotor system in which commands for stabilizing gaze are generated through integration on multiple, distributed timescales.

## INTRODUCTION

Motor plants transform commands from motor neurons into action. Because motor plant responses to neural drive can be complex and history-dependent, commands needed to elicit a particular movement differ from the desired patterns of muscle activation. Standard models of motor control assume that premotor circuitry generates appropriate motor commands by filtering intended muscle activation through an inverse model of the plant being controlled (Kawato 1999; Lisberger 2009). This filtering is then cancelled by the plant’s response to the command, resulting in the intended movement. In this manner, the filtering via the inverse model compensates for the response properties of the plant.

This inverse model framework has proven useful in understanding motor command generation in the oculomotor system (Figure 1A; Green et al. 2007; Robinson 1989; Van Opstal et al. 1985). In the classical view, based on the work of Robinson (1964), passive oculomotor plant behaviour in the horizontal plane is modelled by a pair of one-dimensional viscoelastic (Voigt) elements in series, each characterized by a time constant that dictates the exponential time course of the element’s length change following a step change in applied force. These time constants have been estimated to be relatively short: 10-60 ms and 250-660 ms (Goldstein 1984; Optican and Miles 1985; Robinson 1964; Sklavos et al. 2006; Stahl and Simpson 1995; Stahl et al. 2015). The motor command required for a step change in eye position (saccade), determined by inverting this two-element plant model, is composed of three components, each of which compensates for a different aspect of the plant model’s response properties (Goldstein 1984; Optican and Miles 1985; Robinson 1964). The first component is a brief eye velocity-encoding burst of firing (termed the “pulse”) that overcomes plant viscosity to quickly pull the eye to a new position. The second is an eye position-encoding “step” that counters the plant’s elasticity to maintain the eye at a fixed position during fixation. The final component is an exponential decay (termed the “slide”) that reflects the attenuating force needed to stabilize gaze as the viscoelastic elements equilibrate following the saccade.

**Figure 1.**
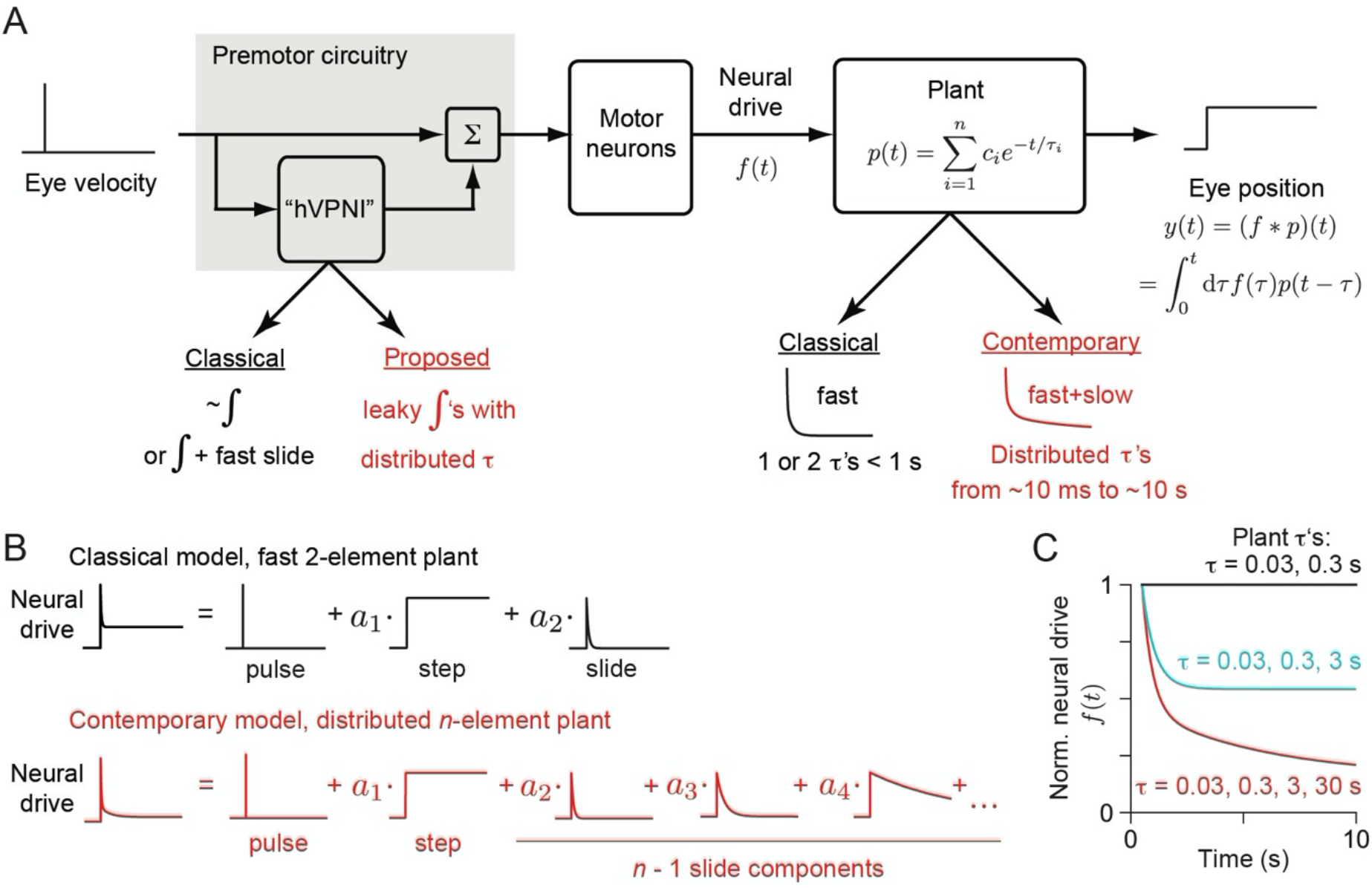
Models of gaze stabilization. (A) Schematic illustration of the conversion of eye velocity commands into the neural drive *f*(*t*) necessary to maintain stable fixation. This neural drive is conveyed by the ocular motor neurons to determine eye position *y*(*t*). The oculomotor plant is modelled as a linear filter *p*(*t*) operating on this drive. Premotor circuitry generates components of *f*(*t*) from signals encoding eye velocity. The classical view, which predates recent results, held that the plant can be characterized by components that relaxed on 10’s and 100’s of ms timescales. Gaze stability on longer timescales then would require the generation of a neural drive that approximates the temporal integral of eye velocity. Here we argue that evidence for a broad distribution of response timescales in the plant redefines the role of premotor circuitry as involving a summation of leaky integrations on distributed timescales. hVPNI: horizontal velocity-to-position neural integrator. (B) Decomposition of the neural drive needed to stabilize gaze under classical and contemporary models of the oculomotor plant. (C) Neural drive required to maintain stable fixation, from 500 ms to 10 s after saccade termination, for a classical plant model (black) and ones with longer response timescales (cyan, red).

Each of these components appears to be reflected in the firing patterns of ocular motor neurons during horizontal eye movements (Robinson 1981). The pulse is conveyed by saccadic burst neurons that project to the ocular motor nuclei. The step component has been observed in the firing of premotor neurons constituting the velocity-to-position neural integrator for horizontal eye movements (hVPNI), which receives eye velocity-encoding bursts and appears to compute their temporal integral, producing the step (Figure 1; Cohen and Komatsuzaki 1972; Robinson 1989; Scudder et al. 2002; Skavenski and Robinson 1973). The slide component has also been observed in hVPNI neurons (Aksay et al. 2000; McFarland and Fuchs 1992); its origin is less clear, but may reflect an imperfect integral of the burst inputs that “leaks” away over time with a particular time constant.

More recent work has suggested a need for substantial modifications to the classical view of eye movement command generation based on a two-element plant model. Sklavos et al. (2006; 2005) found evidence of additional time constants on the order of 1 and 10 s following long steps of force externally applied to the eye. Quaia et al. (2009) analysed the response of primate extraocular muscle to elongation steps, demonstrating muscle tension relaxation on a wide range of timescales, with time constants ranging up to at least 40 s. Davis-Lopez de Carrizosa et al. (2011) measured lateral rectus muscle tension in cats, finding that it decays on timescales of 1 to 10 s during fixations between saccades while eye position is approximately stable. Additionally, firing rates of abducens motor neurons in primates during different types of eye movement are not consistent with a common two-element plant model (Sylvestre and Cullen 1999). These results support an expanded model of the oculomotor plant having several viscoelastic elements with time constants distributed across several orders of magnitude, from 10 ms to 10 s (Sklavos et al. 2005, 2006). The long timescale responses of such plant models imply that additional drive components that decay on long timescales (> 1 s) are necessary to produce stable fixations (Figure 1B,C; Sklavos et al. 2005).

Nevertheless, questions remain about the relevance of long response timescales in the awake, behaving (active) state. Previous measurements of oculomotor plant responses in alert animals have been limited to short time intervals (< 400 ms in monkey, Anderson et al. 2009; < 2 s in mouse, Stahl et al. 2015), precluding observation of long timescale responses. These studies also used brief eye position steps < 1 s, which would not appreciably deform viscoelastic elements with long time constants (Anderson et al. 2009; Sklavos et al. 2006). Furthermore, previous fitting of oculomotor plant models has depended on the assumption that viscoelastic elements were at equilibrium (Sklavos et al., 2005) and was therefore not suitable for fitting model parameters in awake animals.

In addition, questions remain about the relation between long response timescales in the plant and firing dynamics in the hVPNI. We have reported that during fixations, firing rates in hVPNI neurons do not simply encode eye position and a single fast slide, as predicted by classical models, but instead decay on long timescales that vary across an order of magnitude within individual larval zebrafish (Miri et al. 2011a). Such heterogeneity in firing rate decay timescales has also been measured during fixation in adult goldfish hVPNI (Miri et al. 2011a) and in monkey oculomotor integrator neurons (Joshua et al. 2013). In cats, abducens firing after saccades decays on varying timescales generally greater than 1 s, with such decays believed to arise from the oculomotor integrator (Davis-Lopez de Carrizosa et al. 2011). While heterogeneous firing rate decays may reflect an inverse model that helps stabilize a plant with distributed response timescales, it remains to be seen whether firing in the hVPNI could constitute a signal sufficient to do so.

Here we used measurements of oculomotor plant dynamics and analysis of previously obtained neural recordings in the larval zebrafish to assess whether the hVPNI implements an approximate inverse model that accommodates long timescale behaviour of the oculomotor plant to promote gaze stability. In both anaesthetized and awake animals, we performed mechanical displacements sufficiently long enough (~10–100 s) to appreciably deform elements with long timescale responses. We developed methods for analysing the eye’s return from long displacements in the active state. Our measurements demonstrate the existence of both short (< 1 s) and long response timescales in the larval zebrafish oculomotor plant in both the anaesthetized and active states. We used these measurements to fit oculomotor plant models for each larva, employing a new method that does not require an equilibrium assumption. Using these plant models, we then compared the predictions of the neural drive during active state fixations with previous measurements of activity in the larval zebrafish hVPNI. Analysis of the distribution of decay times seen in hVPNI neuron firing rates suggests that this distribution is sufficient to enable stabilization of an oculomotor plant with the distributed response timescales we observed. Our results support a view of integration in the oculomotor system in which hVPNI firing, rather than purely or primarily encoding eye position, compensates for plant viscoelasticity by generating firing rate decay on multiple, distributed timescales (Figure 1).

## METHODS

### Ethical approval

All experiments were performed in compliance with protocols approved by the Princeton University Institutional Animal Care and Use Committee (protocols #1726 and 1863), and in accordance with the policies of the *Journal of Physiology.* Following these guidelines ensured that animal distress was minimized in the course of our study.

### Framework for modelling oculomotor plant

We modelled the oculomotor plant as a combination of viscoelastic Voigt elements in series that respond over *n* effective timescales. This was represented as a linear filter whose impulse response function consists of a sum of exponentially decaying components (Robinson 1964; Sklavos et al. 2005),

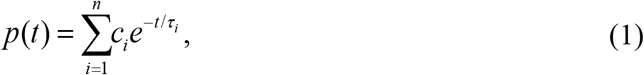

where *c_i_* >0 is the coefficient of the component with time constant *τ_i_*. The time constants can be identified by finding the step response of the system. If a force *f*(*f*) is applied to the system at time *t* = 0, the measured eye position *y*(*t*) will be a convolution of the force profile and the impulse response,

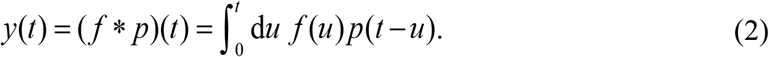

If the applied force is stopped at time *t* = *t*_0_, then the measured eye position at later times will be

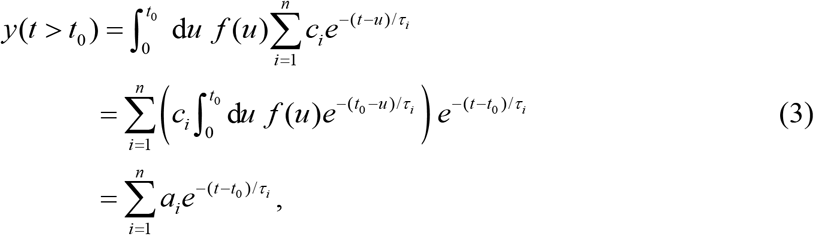

that is, the response after release will be a sum of exponentials with the same time constants as the plant.

Based on the above, in order to model the oculomotor plant, we took the following steps, detailed in the sections below:

1. Apply a transient external force (“displacement”) resulting in a step change in eye position, and measure the eye position after release from displacement (“step response”).
2. Fit a multiexponential model to the step response and extract the plant time constants {*τ_i_*}.
3. Find the plant coefficients {*c_i_*} by fitting a model of the form in equation (2) to the eye position during and after displacement.

### Step response measurement

*mitfa^-/-^* (nacre) mutant zebrafish (*Danio rerio*) larvae (Lister et al. 1999) ages 5-8 days post-fertilization were used for all experiments. We obtained the nacre strain from Zebrafish International Resource Center. Embryos were reared in egg water (Westerfield, 2007) in petri dishes in an incubator at 28°C on a 12 h light/12 h dark cycle. Larvae at this age feed from their yolk and additional food was not provided.

To enable eye tracking, larvae were immobilised by embedding in a thin layer of 1.7% low melting point agarose (SeaPlaque, Lonza) immediately prior to data collection, and the agarose was removed from around the eyes to allow free eye movement. A rectangular agarose block containing the larva was excised and mounted on a Sylgard platform in a water-filled chamber. Individual larvae were embedded for no more than 3 hours. Following data collection, larvae were removed from agarose and immediately euthanized by submerging in ice water for > 5 minutes, to which bleach was added to a concentration of 1% by volume.

We used one of two methods to anaesthetize larvae. Ethyl 3-aminobenzoate methanesulfonate (MS-222, Sigma; *n* = 10 larvae) was gradually added to the chamber water to achieve a concentration at which spontaneous eye movements stopped. Final concentrations were between 0.005 and 0.015% (weight/volume). For ketamine experiments (*n* = 6 larvae), embedded larvae were incubated for 15 minutes in 0.5% (weight/volume) ketamine prior to mounting in the chamber; no ketamine was added to the chamber water. This ketamine concentration was found to be sufficient to abolish spontaneous eye movement in most larvae. Data were not collected from larvae that performed eye movements under anaesthesia. Separate sets of larvae were used for each of the experimental groups: MS-222 anesthetized, ketamine anesthetized, and awake.

The left eye’s response to step displacement was measured in the dark. In order to displace the eye, a blunt probe (hemispherical tip ~30 μm in diameter) controlled by a hydraulic micromanipulator (Siskiyou) was brought toward a point ~50 μm temporal of the centre of the left eye at an oblique angle relative to the eye’s minor axis (considering the eye as an ellipsoid; Figure 2A). After gently contacting the eye, the probe was advanced to rotate the eye temporally in the horizontal plane. Probe advancement was performed quickly, taking less than a second. For active state measurements (*n* = 6 larvae), this displacement was performed 5-7 s following a saccade in which the eye moved nasally. This allowed the expected position of the eye in the absence of the displacement to be estimated by extrapolating a fit to the eye position between the saccade and the displacement (see below). The eye was released by quickly retracting the probe. Eye position was tracked at ~70 Hz for > 60 seconds following release using methods previously described (Beck et al. 2004; Miri et al. 2011b). Briefly, a 945-nm LED illuminated the chamber from below while a mirror, long-pass filter, and charge-coupled device (CCD) camera above the chamber collected video images that were processed in real-time to extract eye position measurements using software custom written in LabView (National Instruments). In this software, two regions of interest (ROIs) that included either the eyes or a fixed segment of the body were drawn on a reference CCD image. During data collection, the ROIs were thresholded, the two largest objects within the eye ROI were defined as the eyes, the largest object within the body field was defined as the body segment, and the edges of these three objects were smoothed. The body axis was defined as the line connecting the centre of the body segment and the midpoint between the centroids of the eyes. Horizontal eye positions were measured as the angle formed by the major axis of the eye and the body axis. Eye position measurements were digitized by a Digidata 1440A (Molecular Devices) and recorded at 5000 Hz in Clampex (v.10, Molecular Devices). Visual inspection of eye tracking images indicated that eye shape was at most minimally disturbed by contact with the probe and any disturbance was confined to the site of probe contact. Contact with the probe did not cause visible deformation of the eye that extended appreciably into the displacement response.

**Figure 2.**
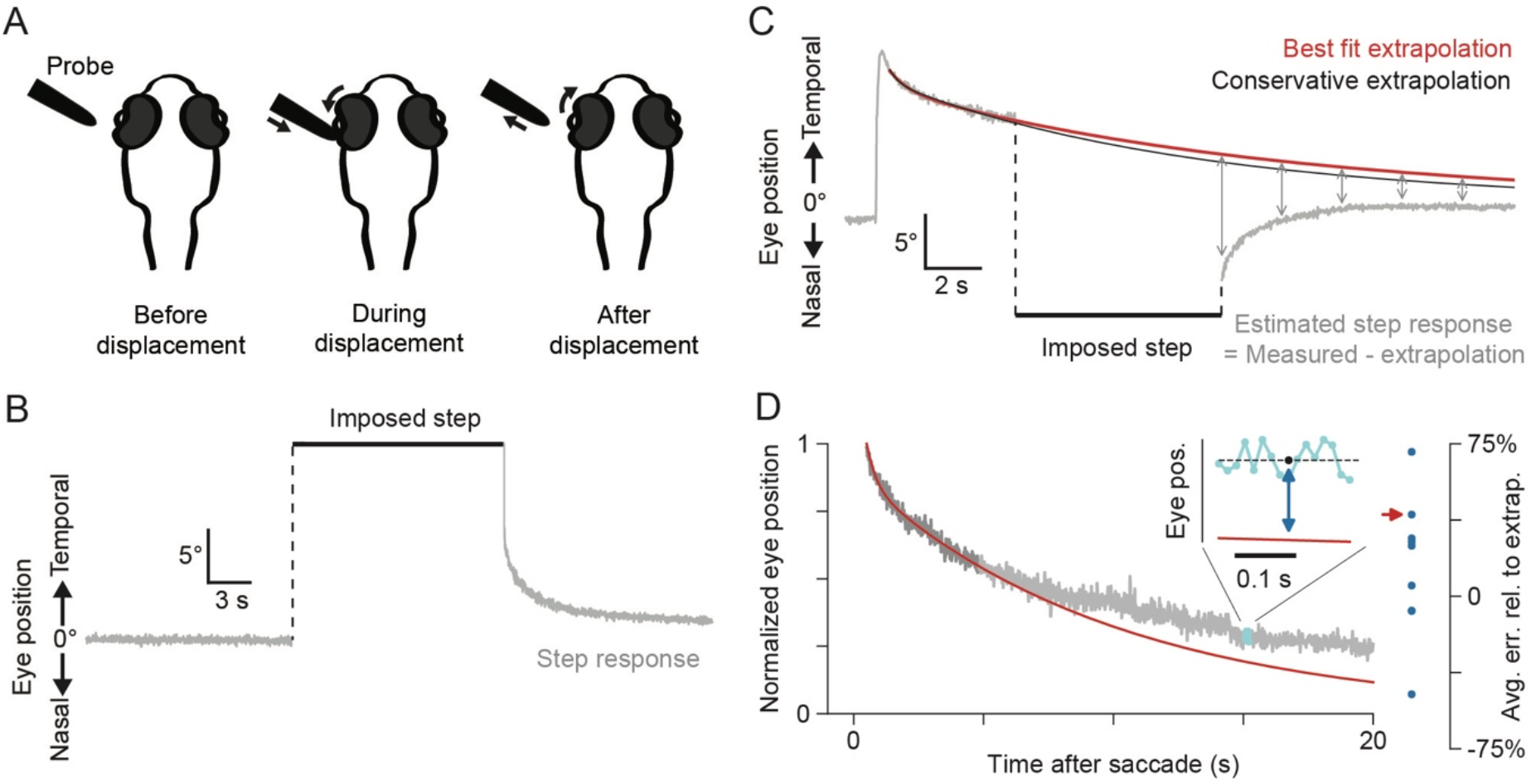
Methodological approach. (A) Schematic illustration of the method used to measure the oculomotor plant step response. A blunt probe controlled by a hydraulic micromanipulator was used to transiently displace the eye. (B) In anaesthetized larvae, the step response was the eye position measured after release from an imposed step displacement. (C) In awake larvae, the eye after displacement was assumed to return to the position it would have occupied had no displacement occurred. Thus, the step response was estimated to be the measured eye position after release from the step displacement (grey), minus the extrapolation of pre-displacement eye position (red). A faster decaying conservative extrapolation (black) was also calculated. (D) Left: to validate the extrapolations, fits to the initial portion (darker grey) of long unperturbed fixations were extrapolated (red), and the fractional error of the extrapolation relative to the true eye position was averaged over a ~230 ms window centred at 14.64 s (inset: cyan, data; dashed black, mean of these data). Right: the average relative error across the 230 ms window for each measured fixation (red arrow corresponds to the fixation shown on the left).

In each anaesthetized larva, two displacements of differing duration were performed: 10 and 60 s in MS-222 experiments, 15 and 90 s in ketamine experiments. Displacements ranged from 14.8 to 22.2° under MS-222, and 8.5 to 20.9° under ketamine. Pairs of displacements performed on individual larvae were nearly equal, differing on average by only 1.5°. In awake larvae, up to five displacements were performed on each larva, each between 6.5 and 8.5 s in duration, and 11.6 and 26.0° in size. Active state trials in which spontaneous eye movements occurred during the applied displacement or within 8 s following the release were discarded. As a result, at most two responses from each larva were analysed. Spontaneous eye movements during displacement could be identified by motion of the undisplaced eye. At least 10 minutes elapsed between displacements.

### Step response fitting

Eye position during the displacement prior to release was measured from an image captured while the eye was displaced. The time of release was defined as the time of the last sample during which the probe was contiguous with the eye in the video image. For MS-222 and ketamine experiments, baseline eye position was measured as the mean eye position during a 50 second epoch preceding the displacement and was subtracted from the eye position time series.

For active state responses, the centre of gaze was estimated from a plot of eye velocity versus eye position (a “PV plot”; Becker and Klein 1973; Goldman et al. 2002) assembled as follows from at least three minutes of eye position data collected during spontaneous eye movement prior to the displacement. First, data from 100 ms prior to each saccade until 500 ms after each saccade were discarded. The remaining time series data were divided into non-overlapping 0.3 s segments. Next, the mean eye position and the slope of a least-squares fit line to eye position over each segment were calculated to define the two coordinates of each point for the PV plot. Finally, a linear function was least-squares fit to these points. The intercept of this function with the eye position axis was defined as the centre of gaze and subtracted from all measurements in the eye position time series.

Since data were initially acquired at ~70 Hz and digitized at 5000 Hz, we downsampled eye position time series before performing any data analysis. We therefore subsampled traces every 72 time points, resulting in new traces at 69.44 Hz that we then analysed. Subsequent analysis of the power spectra of eye position traces showed anomalously large peaks at ~30 Hz, which were likely artefacts. These were removed, while preserving the phase in each frequency bin, by scaling the amplitudes of the Fourier transform in the peaks so that the amplitude was equal to that of the mean amplitude of the 6-10 frequency bins surrounding the peak (3-5 closest on each side of the peak, chosen by visual inspection of power spectrum). All analyses were performed in Python 3.7, using the Scientific Python stack (SciPy and NumPy).

#### Anaesthetized step responses

Eye position step responses (Figure 2B) in anaesthetized larvae were fit with a multiexponential model,

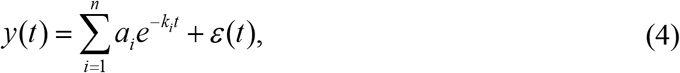

where *ε* is independent Gaussian noise with mean 0 and variance *σ*^2^, *k_i_* = 1/*τ_i_* are inverse time constants, and the number of components *n* ranged between 1 and 6. Each component amplitude was constrained to be nonnegative, *α_i_* ≥ 0. Eye positions were normalized to be 1 at the time of release, and accordingly the sum of the component amplitudes in the model was constrained to equal 1. This allowed for comparison of step response fits even though displacement amplitudes varied. Visual inspection of these time series near the release time found ringing/oscillation present during the initial 50-200 ms following release in some cases, perhaps resulting from the manual control of the hydraulic manipulator. We therefore analysed responses beginning 230 ms and ending 60 s after release time.

For each larva, we simultaneously fit the short and long step response with models of the form in equation (4) for each value of *n* between 1 and 6. For each value of *n*, we defined the best *n*-component model to be the one which maximized the sum 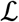 of log-likelihoods 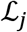 for each response *j*,

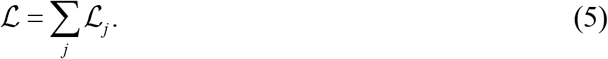

Here,

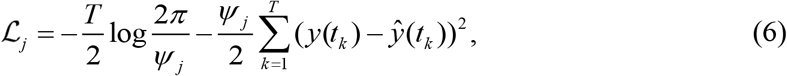

where *T* is the total number of time points {*t_k_*} recorded in the response, *y_j_* is the recorded eye position, *ŷ_j_* is a model in the form of equation (4), and, for numerical stability, we defined 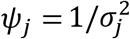 where *σ_j_* is the standard deviation of the noise. We allowed the fits to each of the two response durations to have different sets of component amplitudes {*a_i_*}, but we required a single set of inverse time constants [*k_i_*], with corresponding time constant values constrained to be greater than 43.2 ms, or 3 samples. Separate *σ_j_* were fit for each response. To find sets of parameters that maximized 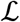, we used the nonlinear solver Truncated Newton Conjugate-Gradient (TNC; implemented in the “optimize” library of SciPy), which we provided with an analytical formula for the gradient of 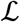 that was derived by hand. We used 100 initial sets of parameter values by choosing component amplitudes uniformly at random between 0 and 1 and then dividing each coefficient by the sum of all component amplitudes. Initial time constants were chosen by taking random powers of 10, generated by first taking *n* evenly spaced powers between −1 and 2, and adding Gaussian noise with mean 0 and standard deviation 0.1 to each. By examining the mean squared error curves of fits as *n* increased, we saw clear “elbows” after which fit quality stopped visibly improving. Separately for each larva, we called the value of *n* at which this elbow occurred *n*\*. For the sake of parsimony, we picked the best overall model for each larva to be the best *n\**-component model.

To examine the sensitivity of the parameter estimates, we used a parametric bootstrap procedure as follows. For each larva, we used the best step response fit (component amplitudes, time constants, and measurement noise variances) to generate 100 new eye position traces according to the model in equation (4), and re-ran our fitting procedure on each of these, then calculated the standard deviations of the resulting bootstrap parameters, which we used as an estimate of the standard deviations of the true parameter distributions.

In order to determine the necessity of including long time constant components, we next repeated the above procedure to find the best fits for each larva when time constants were constrained to be less than 10 s.

#### Active state step responses

Because the eye moves spontaneously in the active state and is not at the centre of gaze prior to the imposed displacement, we did not model the active state step response as a decay toward the centre of gaze. Rather, we modelled it as returning to where the eye would have been had the displacement not occurred (Figure 2C). Let *y*_target_(*t*) be the position of the eye if no displacement had occurred. Then, we modelled the active state step response as

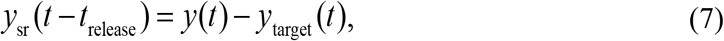

where, as before, *y*(*t*) is the measured eye position.

We determined *y*_target_(*t*) by extrapolating the changing position of the eye, based on the 4-6 s of post-saccadic eye relaxation immediately preceding displacement (Figure 2C). We found that this relaxation could also be modelled well by equation (4), which we fit by maximum likelihood estimation to the eye position from 500 ms after the previous saccade to displacement onset, using the method described above for anaesthetized step responses. For all but one active state response we picked a 2-component model for the best extrapolation, as there was negligible improvement in fit quality for >2 components. For the remaining response, we used a 3-component model.

To evaluate the quality of this extrapolation technique, we identified long nasal fixations lasting at least 15.26 s in eye position recordings prior to each active step response. We then performed the same fitting procedure used to find *y*_target_(*t*) above, here fitting eye position from 500 ms to 4.8 s post-saccade (Figure 2D). We then calculated how much the resulting fit function deviated from the true eye position in a ~230 ms window of time centred at 15.16 s post-saccade by calculating the error relative to the extrapolation at each data point in this interval,

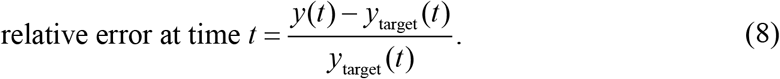

For each fixation, we then calculated the average relative error over the 16 data points in this window (Figure 2D, inset; black point indicates the centre point of this window at time *t*_extrap_ = 15.16 s). A negative relative error meant that the extrapolation decayed more slowly than the true fixation. We found that the average relative error was positive, so that fits were likely to decay more quickly than the true fixation, but with relatively large variance (Figure 2D, right; mean ± standard deviation [sd] = 0.16 ± 0.33; *n* = 9). 3 out of the 9 fixations resulted in a negative average relative error, where the extrapolated eye position decayed more slowly than true eye position. Thus, a step response estimated from these extrapolations would also decay more slowly than the “true” step response and would be more likely to have long time constants. In order to make our fitting procedure more robust to such an artefact, which could lead to spurious identification of long time constant decays, we sought to also fit a more quickly decaying, “conservative” extrapolation of eye position for each active state step response.

We fit the conservative extrapolations by finding fit functions which fit the first 4-6 s of eye position well, but also decayed faster than the best extrapolation. We did this by including a penalty term in our optimization that incentivized fit functions to pass through a virtual point at time *t*_extrap_ that was closer to the null eye position than the best extrapolation would be. First, we defined Δ to be the most negative relative error, averaged over the window described above (Figure 2D, right, bottom blue point). Then, starting with the best extrapolation (Figure 2C,D, red curve), we fit a higher order model, i.e., *n* + 1 components if the best extrapolation needed *n* components, by maximum likelihood estimation, but where we augmented the model (4) by fitting an additional point *y*(*t*_extrap_) = (1 + Δ)*y*_target_(*t*_extrap_). All of the fixations before displacement were less than 7 s long, so this additional point did not overwrite any actual data points. We let the noise at this additional point, *ε*(*t*_extrap_), be normally distributed with mean 0 and variance *σ/λ*. Increasing *λ* increasingly penalizes fits that do not pass through *y*(*t*_extrap_). By bisection search on *λ*, we allowed the procedure to choose a fit whose sum of squared errors over the real data deviated from that of the best extrapolation by ~10% as follows (Figure 2C,D, black curve). For each value of *λ*, we performed maximum likelihood estimation from 100 starting points and picked the most parsimonious model, as described above (Anaesthetized step responses), to perform this comparison. The bisection search terminated when the deviation of the sum of squared errors fell between 9.9 and 10.1%. The eye position predicted by the maximum likelihood model for the value of *λ* for which the bisection search terminates was defined to be the conservative extrapolation.

For both the best extrapolation and conservative extrapolation, we calculated the corresponding step response time series *y*_sr_(*t*). For each displacement in each larva, we fit multiexponential models to both of the resulting active step response time series from 230 ms to between 8 and 14 s after release from displacement, with the fits performed as above for anaesthetized responses. After discarding responses due to spontaneous eye movements, a single response was fit for 3 of 6 larvae and a pair of responses was fit for the remaining 3. We again ran the parametric bootstrap procedure, as in the anaesthetized case above, here using the best three-component fits in order to evaluate the sensitivity of parameter estimates. In order to evaluate the necessity of long time constant components, we also fit responses while constraining the time constants to be less than 5 s. This was again done assuming the best and conservative extrapolations.

For comparison of anaesthetized and active state results, we also fit three-component models as above to just the first 15 s of eye position following the 10 s displacement in larvae anaesthetized with MS-222.

### Oculomotor plant model estimation

To estimate an oculomotor plant model for each larva, we used measured step responses to calculate parameters of the linear filter given by equation (2) above. We assumed that each step response was the result of an applied force convolved with a linear filter representing the oculomotor plant. As described above, a linear filter model implies that the eye position after release from displacement will have the same number of components and the same time constants as the plant (see equation (3)). However, the coefficients of the plant {*c_i_*} must still be determined. Because the profile of the applied force is unknown, finding the coefficients of the plant requires simultaneously finding the appropriate applied force profile. This is a “blind deconvolution” problem and is generally under-constrained. Here, however, we have two important constraints that facilitate finding a solution: first, we partially know the applied force profile, i.e., that it is zero after the time of release; and second, the plant is assumed to be a sum of exponentials with known time constants.

For each larva, we assembled eye position time series *y_j_* starting from the onset of displacement for each response *j* until 8 to 14 s after release. All the anaesthetized larvae had 2 responses each, 3 of 6 awake larvae had two responses, and the remaining three had a single response. Eye position was considered to be constant during displacement (see Step response fitting). For awake larva time series, we used estimated step responses *y*_sr_(*t*) resulting from both best and conservative extrapolations, as described above. Similar to the fitting of step responses, we removed the first 230 ms of data after release from displacement due to ringing/oscillation shortly after release. Data points in this epoch were replaced with the prediction of the best fit exponential model to the step response.

For linear filter estimation, we then defined the following loss function,

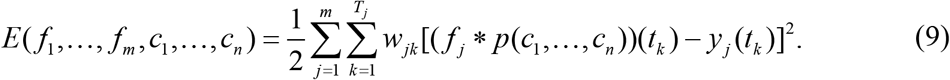

Here, *m* is the number of responses for the larva, *f_j_*(*t*) is the time-varying force applied for response *j*, *p*(*t*; *c*_1_,…,*c_n_*) represents the plant model parameterized by variable coefficients {*c_i_*} and fixed time constants {*τ_i_*}, time is discretized as *t_k_* = *k*Δ*t* where 1/Δ*t* is the sampling rate, and *T_j_* is the number of time points in the response *y_j_*. Each data point in the loss function is weighted by a factor

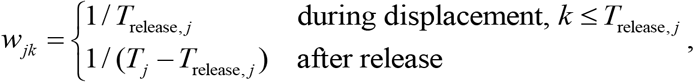

where *T*_release,*j*_ is the number of data points in the displacement period. This weighting causes the loss function to be a sum of the mean squared fit error during displacement and the mean squared fit error during the step response, with errors in these periods weighted equally even though the period lengths are heterogeneous. We approximated convolution of continuous signals with a discrete convolution, since Δ*t* is small,

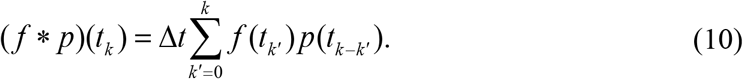

The blind deconvolution can then be computed by finding the forces {*f_j_*} and plant coefficients {*c_i_*} such that the loss *E* is minimized. First, we used the nonlinear solver TNC from 100 starting points that consisted of {*c*_*i*,initial_} chosen uniformly at random on (0, 1) and normalized to sum to 1. These preliminary results were used as the initial point, 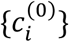 for an alternating least squares procedure, as follows:

1. *Applied force estimation step.* On iteration *l* + 1, given estimates for the plant coefficients 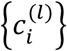, solve the optimization problem

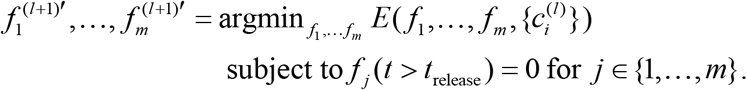
2. *Plant estimation step.* Given these estimates of the applied forces, update the estimate of the plant coefficients,

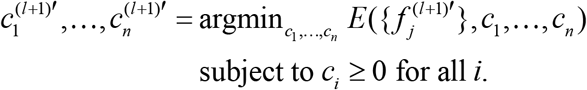 Note that the solutions obtained by this procedure are not unique: for example, if all the plant coefficients are divided by a constant *K*, then as long as the resulting applied forces are also multiplied by *K*, the new scaled coefficients still solve the problem. We resolve this degeneracy by imposing the condition that the final plant coefficient estimates sum to 1:

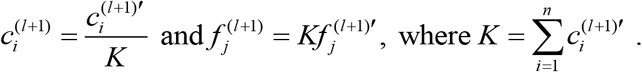

These two steps are then repeated until the decrease in error is smaller than a threshold, *E*^(*l*+l)^ – *E*^(*l*)^ ≤ 10^-8^. Note that the error cannot increase on any step, because in the worst case the same solution as the previous iteration can be chosen. Because the discrete convolution given by equation (10) is linear, we can write it as a matrix-vector product, and each of the two steps above can be solved efficiently and accurately by a linear least squares solver (lsq_linear in SciPy’s “optimize” library). The plant model for each larva was chosen to be the one of the 100 solutions that had the smallest final value for *E*. We repeated this procedure using time constants and eye positions resulting from the conservative extrapolations in awake larvae.

For each plant model, we calculated the fractional contribution of each exponential component to the total area under the curve by integrating over the range *t* = 0 to *t* = 60 s for anaesthetized plants, and to *t* = 20 s for the active state plant models.

### Neural drive estimation

Using the active state plant models, we estimated the force needed to produce the eye position profile observed during fixation. We assumed this force was proportional to the neural drive output by motor neurons. We computed neural drive estimates by deconvolving active state eye position during the period of fixation preceding displacement with the corresponding plant model for each larva, and separately for a classical two-element plant model with time constants of 20 and 200 ms and coefficients of 0.4 and 0.6, respectively (Robinson et al. 1990). In order to reduce noise, we used *y*_target_(*t*) as a smoothed representation of eye position before displacement (see Step response fitting), and performed the deconvolution by solving a linear least squares problem with non-negative least squares, as described above (Oculomotor plant model estimation). We measured the persistence of both the estimated neural drive and the corresponding eye position by summing the time series during fixation starting from and normalized to the value at 0.25 s post-saccade, and dividing this sum by the number of elements in the time series (Lee et al. 2015). With this measure, a perfectly stable time series would have a persistence value of 1.

We also directly calculated the time constants of the slide components of the neural drive for each plant by calculating the poles of the Laplace transform of the neural drive. This was equivalent to finding the roots of the polynomial

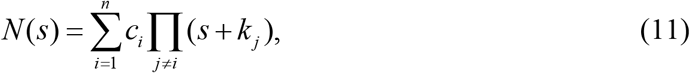

where *c_i_* and *k_i_* are the coefficient and inverse time constant of the *i*th plant component, as in equations (1) and (4) (see Mathematical Appendix, Analytical calculation of neural drive for a multiexponential plant, for derivation). This was done numerically using the function “roots” in NumPy.

### Comparison of estimated neural drive to hVPNI activity

#### Optical recordings

To compare the required neural drive to the activities of cells in the hVPNI, we used previous optical recordings of somatic calcium-sensitive fluorescence in a separate set of 6 larvae to estimate neuronal firing rates during fixations (Miri et al. 2011b). Calcium-sensitive dye loading and optical recording methods are described in the original reference. Data were collected using Oregon Green BAPTA-1 AM (Invitrogen) on a custom-built laser-scanning two-photon microscope that allowed synchronous eye tracking and fluorescence image time series collection from sagittal planes within the hindbrain. Fluorescence data acquisition and microscope control were performed using Cfnt v.1.529 (Michael Mueller, MPI, Heidelberg). Images were 256 x 256 pixels spanning 100 μm x 100 μm regions and acquired at 2 ms per line (~2 Hz) in time series of 750 frames. For each larva, five or six fluorescence image time series were collected from image windows lying in parasagittal planes at fixed dorsoventral and rostrocaudal coordinates in the ventral ~2/3 of the caudal hindbrain (rhombomere 7/8). All data analysed here were collected in the dark to eliminate visual feedback.

#### hVPNI firing rate estimation

For each cell, we determined baseline fluorescence to be the smaller of two quantities: the mean of the saccade-triggered average fluorescence from 2 s to 1 s before ipsiversive saccades, and the mean of the saccade-triggered average fluorescence from 4 s to 5 s after contraversive saccades. We fit a calcium impulse response function (CIRF), modelled by a single exponential decay, to baseline-subtracted contraversive saccade-triggered average fluorescence time series, as in previous studies (Daie et al. 2015; Miri et al. 2011b) and calculated the coefficient of determination, R^2^, for each fit. As in previous studies (Miri et al. 2011a, Miri et al. 2011b), cells for which CIRF fits had R^2^ ≥ 0.5, and the Pearson correlation between saccade-triggered average fluorescence and CIRF-convolved eye position after a contraversive saccade was > 0.5, were used (166 of 195 cells).

For each cell *j* satisfying these criteria, we modelled baseline-subtracted ipsiversive saccade-triggered average fluorescence *x_j_* as a convolution between the CIRF and the sum of two components: a delta function modelling the saccadic burst and a multiexponential function representing the saccade-triggered average firing rate,

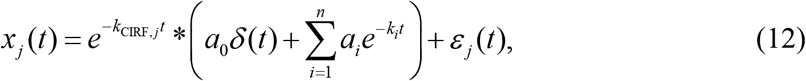

where *k*_CIRF,*j*_ is the cell-specific inverse CIRF time constant, *k_i_* are inverse time constants of the saccade-triggered average firing rate, with coefficients *a_i_* ≥ 0 for all components *i*, and *ε_j_* is Gaussian noise with mean 0 and variance 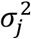. For each number of exponential components *n* between 1 and 3, we solved for the coefficients {*a_i_*} and inverse time constants {*k_i_*} that minimized the mean squared error of the model fit to data, using the nonlinear solver TNC from 100 initial sets of parameters. For each number of components *n*, we picked the best *n*-component model fit to be the one with the smallest mean squared error. Separately for each cell, we called *n*\* the value of *n* after which adding another component decreased the mean squared error by less than 1%. Then, we defined the best overall firing rate model fit for each cell to be the best *n*\*-component model fit. We analysed responses from cells where the ratio of the sum of squared errors of the best firing rate model fit to the sum of squares of the data (sum of squares ratio, SSr) was less than 0.007 (151 of 167 cells)—cells with SSr greater than this value had fits that were noticeably worse upon visual inspection. For each cell, we calculated firing rate persistence values in the same manner described above for neural drive and eye position persistence (Neural drive estimation).

#### Summary plant and neural drive estimation

For the 9 active state step responses from 6 larvae, we simultaneously fit a single set of time constants, using the same procedure as above (Step response fitting). Then, using these time constants, we performed the blind deconvolution procedure (Oculomotor plant model estimation) to find the best fit plant model for all 9 responses simultaneously (“summary plant”). We then computed the saccade-triggered average eye position using smoothed versions of fixations from each of 6 separate larvae in which optical recording was performed. Smoothing was performed by fitting multiexponential functions to the saccade-triggered average eye position from each animal, as described above (Step response fitting). A neural drive estimate was then generated for each larva by deconvolving the saccade-triggered average eye position with the summary plant.

We then performed a regularized linear regression of the saccade-triggered average firing rates of all hVPNI neurons onto the neural drive estimate for each larva, with regression weights constrained to be nonnegative, using a linear least squares solver (lsq_linear). We used a regularization term equal to the sum of squares (L_2_ norm) of the regression weights of each time series, multiplied by a regularization parameter. We chose this parameter separately for each larva by bisection search so that approximately 50% of the weights were nonzero, broadly in agreement with the fraction of putative hVPNI neurons that synapse onto motor neurons in electron microscopy data (Lee et al. 2015, Vishwanathan et al. 2017).

To show the importance of intermediate timescales in the neural drive (Mathematical Appendix), for each larva we generated 100 synthetic populations of 150 mock cells whose firing rates were each described by an exponential decay. For each population, 75 of the cells’ firing rates had time constants that were randomly drawn from a uniform distribution between 0 and 1 s, and 75 had time constants generated by taking 10 to the power of a uniform random number between 1.3 and 2, resulting in random time constants between 20 s and 100 s. We then performed a regularized linear regression of the mock cells’ firing rates onto the estimated neural drive exactly as for the real cells.

## RESULTS

### Measurement of zebrafish oculomotor plant response under anaesthesia

In order to determine whether the larval zebrafish oculomotor plant, like that of the anaesthetized primate, shows both short and long response timescales (Sklavos et al. 2006; Sklavos et al. 2005), we first measured the response of the eye to horizontal step displacements (“step responses”) of two different durations in larvae anaesthetized with MS-222. Because MS-222 inhibits action potential firing and thus input to neuromuscular junctions, active muscle tone is likely diminished or absent in this state so that the observed responses primarily reflect the passive properties of the plant. Similar amplitude abducting step displacements lasting 10 and 60 s were applied to one eye of anaesthetized larvae (*n* = 10) and the return trajectory of the eye was tracked following release (Figure 2A,B). These experiments and those described in what follows were performed in the dark to prevent the influence of visual feedback. The use of two different step durations helps expose both short and long response timescales; if the plant responded on exclusively short timescales, it would effectively reach steady state within 10 s, so that the responses to 10 and 60 s displacements would be similar.

However, the responses to 10 and 60 s step displacements were strikingly different (Figure 3). To quantify response timescales present in these trajectories, for each larva we simultaneously fit with multiexponential functions the first 60 s following release of responses to both displacements. These functions had from 1 to 6 exponential decay components and were constrained at the time of release to equal the eye position prior to release. For each function with a given number of components, fits assumed a single set of time constants but distinct sets of corresponding amplitudes for the responses to the 10 and 60 s displacements. Oscillatory artefacts sometimes present in responses during the first 200 ms following release, perhaps resulting from the manual control of the hydraulic manipulator, led us to omit the first 230 ms of responses when fitting. Thus, our model fits cannot be expected to well capture response timescales faster than ~100 ms. Single exponential functions failed to capture much response structure, while models with 4 components provided a better fit (Figure 3A,B). Improvements in fit quality as the number of model components increased were similar to those observed previously for plant responses measured in anaesthetized monkeys (Sklavos et al. 2005) and awake mice (Figure 3B; Stahl et al. 2015).

**Figure 3.**
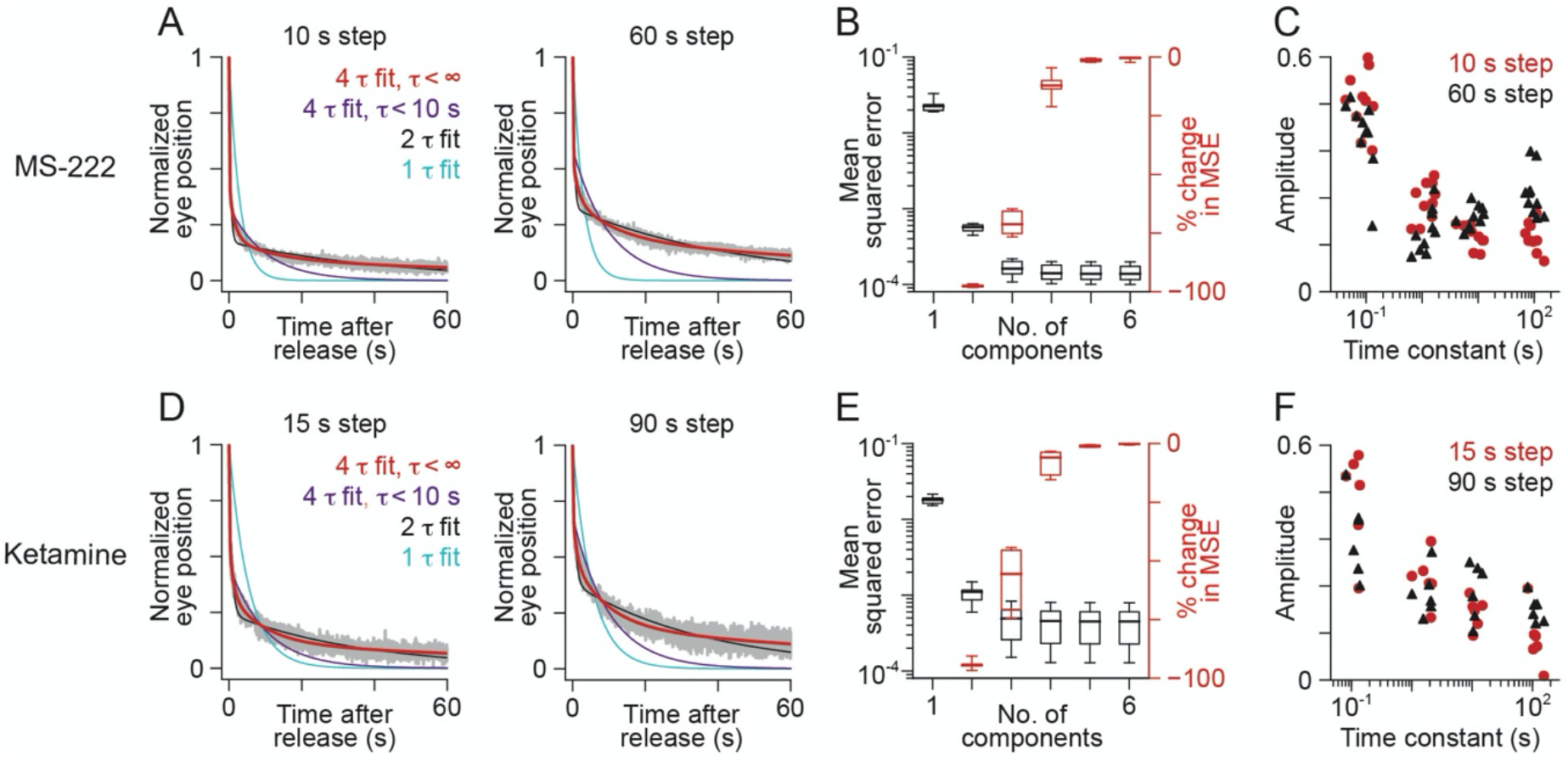
The eye’s return from step displacements in anaesthetized zebrafish larvae reveals short and long response timescales in the oculomotor plant. (A) Example responses of the eye (grey) following step displacements lasting 10 s (left) or 60 s (right), normalized by eye position prior to release, in larvae anaesthetized with MS-222. Coloured traces: simultaneous fits to both displacements with multiexponential models having one (cyan), two (black), or four components with time constants unconstrained (red) or constrained to be less than 10 s (purple). (B) Mean squared error (MSE) of fits with different numbers of components (black) and percent change in MSE compared to the fit with one fewer component (red) in larvae anaesthetized with MS-222. Boxes span the 25th to 75th percentile range; whiskers show maximum and minimum values (*n* = 10 larvae). (C) Component amplitudes and time constants for four-component fits in larvae anaesthetized with MS-222. (D-F) Same as A-C but for larvae anaesthetized with ketamine (*n* = 6), for which step displacements lasted 15 or 90 s.

In order to determine the minimum number of discernible timescales in step responses, we examined the change in mean squared error of fits as the number of exponential decay components used in the fits was increased. For each larva, there was an “elbow” at 4 components, after which the reduction in mean squared error by adding more components was extremely small (Figure 3B). The median values of the time constants of the four-component fits were 0.092, 1.34, 7.95 and 91.6 s (Figure 3C). Bootstrapped standard deviation estimates for time constant values were small relative to the gaps between values for successive components (median sd of estimates ranged between 2.1 and 6.6% as a fraction of the best-fit time constant values). Time constants greater than 10 s were necessary to well fit responses; multiexponential models constrained to have time constants less than 10 s failed to fit step responses as well as unconstrained models having equivalent numbers of components (Figure 3A; 39- to 172-fold increase in MSE for four-component models).

We next measured step responses under another anaesthetic, the NMDA receptor antagonist ketamine. These experiments were performed for three reasons: (a) to demonstrate that aspects of the results obtained with MS-222 are not dependent on the choice of anaesthetic, (b) to better compare our results in the larval zebrafish with those of Sklavos et al. (2005) in the primate, and (c) because there is some indication in mammals that at ketamine doses near the threshold above which the animal becomes unresponsive to eye manipulation, some active muscle tone is preserved, making ketamine anaesthesia potentially closer to the active state (Blanks et al. 1977; King et al. 1978; Sklavos et al. 2005). We did not attempt to verify the presence of active muscle tone under ketamine. After anaesthetizing larvae (*n* = 6) with ketamine, we applied similarly sized abducting step displacements of 15 and 90 s to one eye and tracked its return trajectory following release (Figure 3D).

We simultaneously fit the responses following 15 and 90 s displacements using multiexponential functions as described above. The choice of slightly different step durations here was arbitrary and should not obscure the general agreement this similarity demonstrates. Fit results were similar to those obtained under MS-222 (Figure 3D-F). The median values of the time constants of four-component fits were 0.131, 2.01, 10.7, and 110.2 s (Figure 3F). Bootstrapped standard deviation estimates for time constant values were small relative to the gaps between values for successive components (median sd of estimates ranged between 4.7 and 8.4% as a fraction of the best-fit time constant values). Models constrained to have time constants less than 10 s again failed to fit step responses well (Figure 3D; 8- to 58-fold increase in MSE for four-component models).

Collectively, the data from anaesthetized larvae suggest that the larval zebrafish oculomotor plant, like that of the primate, demonstrates both short (< 1 s) and long (> 1 s) response timescales. Moreover, our observation of time constants spread over several orders of magnitude under both types of anaesthesia matches results from comparable primate experiments (Sklavos et al. 2006; Sklavos et al. 2005).

### Measurement of oculomotor plant responses in the active state

Because active tone in extraocular muscles could influence the response properties of the oculomotor plant, we examined whether long response timescales were discernible in awake, behaving larvae. Previous observations of active state plant responses have been limited to relatively brief time windows (< 400 ms in monkey, Anderson et al. 2009; < 2 s in mouse, Stahl et al. 2015) and relatively short displacements (< 1 s), potentially obscuring the presence of long response timescales. Here we succeeded in measuring active state plant responses of longer duration thanks to the relatively low saccade frequency of larval zebrafish. Five to seven seconds after an adducting saccade in the dark, we applied abducting step displacements lasting 6.5 to 8.5 s. On 9 occasions across 6 larvae, we were able to record responses lasting > 8 s without any interrupting saccades. To our knowledge, active state measurements of comparable duration have not been previously reported.

In order to properly fit these responses, we needed to estimate what the eye position would have been during the responses had the imposed displacements not occurred (Figure 2C). Because eye position decays toward the centre of gaze appreciably during fixations in larval zebrafish, we could not use the eye position immediately prior to displacement as an estimate of the expected eye position in the absence of displacement. Instead, we estimated the expected eye position in the absence of displacement by fitting multiexponential models to eye position between the previous adducting saccade and the displacement, and then extrapolated the model fits forward in time. Models having between 1 and 6 components were initially fit, and the number of components used for subsequent analysis was chosen using the reduction in mean squared error from each added component as described above. In addition to this “best” extrapolated eye position fit, we also performed a second “conservative” fit with faster eye position decay (see Methods, Figure 2D) that conservatively accounts for possible overestimates of expected eye position resulting from extrapolation. We extrapolated both the “best” and “conservative” fit functions through the step and subsequent response epochs to predict eye position had the displacement not been applied. We defined the step response as the difference between the eye position following release and these extrapolated fit functions (Figure 2C, grey arrows).

To quantify response timescales in the active state, we fit multiexponential functions to responses from 230 ms to between 8 and 21 s following release for each larva. For 3 larvae, a single response was fit. For 3 other larvae, responses to two separate displacements were simultaneously fit with a common set of time constants, but distinct component amplitudes as in the anesthetized case. Results assuming the best and conservative extrapolations were generally in agreement. Single exponential functions failed to well-capture response structure (Figure 4A,B). For 5 of 6 larvae, responses were best fit by a three-component model (Figure 4C), while responses from the remaining larva was best fit by a four-component model. Fit improvements upon inclusion of additional components beyond 4 were again generally very small (Figure 4D). Time constants were broadly distributed for all larvae, assuming both the best and conservative eye position extrapolations (Figure 4E,F). Assuming the best extrapolation, the median values of the time constants of three-component fits for all 6 larvae were 0.170, 2.52, and 38.8 s. Bootstrapped standard deviation estimates for time constant values were again small relative to the gaps between values for successive components (median sd of estimates was around 5% as a fraction of the best-fit time constant values). Here again, very long time constants were necessary for an adequate fit; multiexponential models constrained to have time constants no greater than 5 s produced much worse fits to step responses than unconstrained models having equivalent numbers of components (Figure 4A,B; 3- to 459-fold increase in MSE for three-component models using the best extrapolation and 1.3- to 356-fold increase using the conservative extrapolation). Collectively, these results demonstrate that, even when accounting for possible errors in extrapolation, active state step responses also display both short and long response timescales.

**Figure 4.**
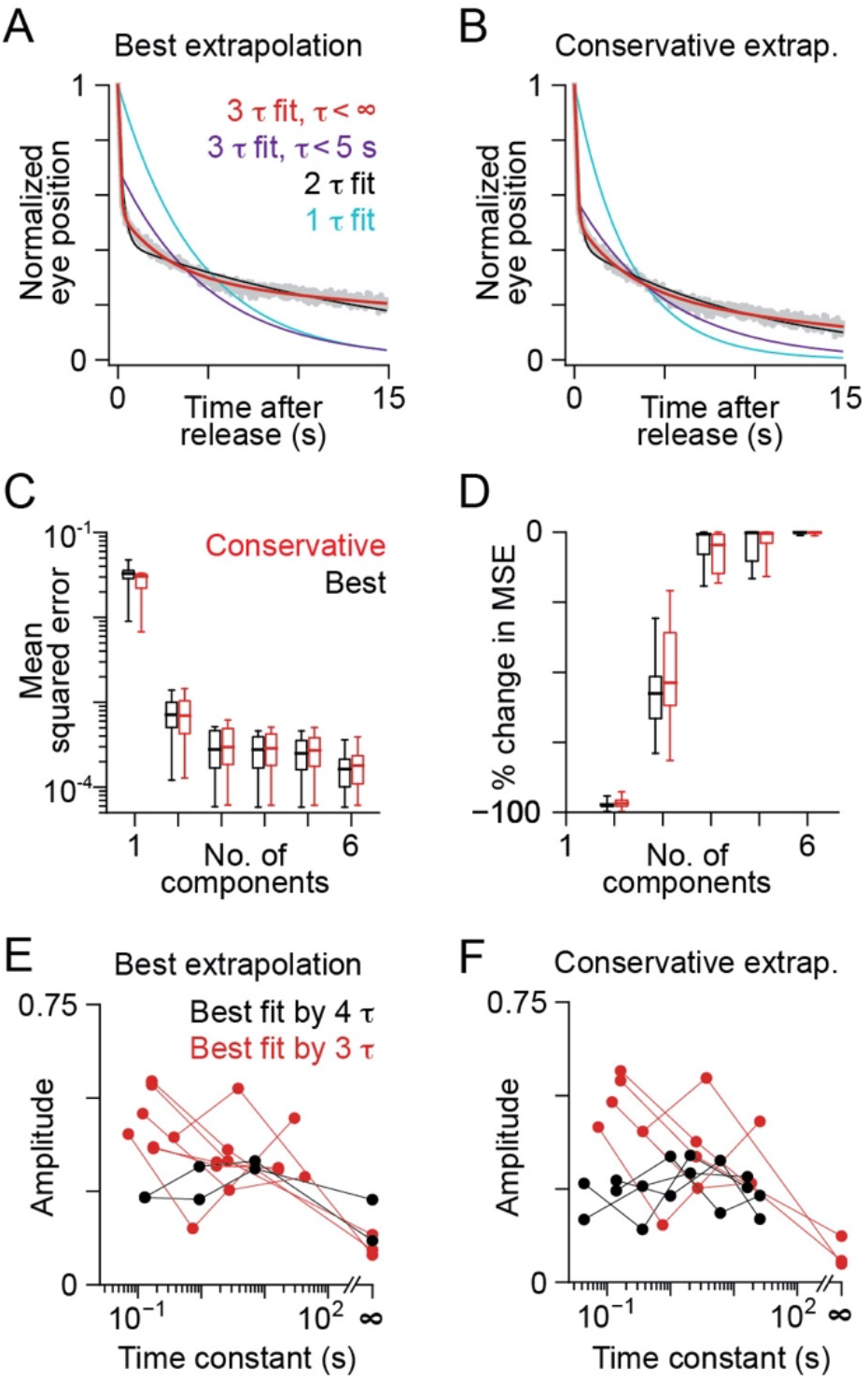
The eye’s return from step displacements in awake zebrafish larvae exhibits short and long response timescales. (A,B) Example active state step response calculated using the best (A) and conservative (B) extrapolation of eye position in the absence of displacement. Fits of multiexponential models having one (cyan), two (black) or three components with time constants unconstrained (red) or constrained to be less than 5 s (purple) are overlaid. (C,D) Mean squared error (MSE) (C) and percent change in MSE (D) for step response fits compared to models with one fewer component, calculated using the best (red) or conservative (black) eye position extrapolations for each larva. Boxes span the 25th to 75th percentile range; whiskers show maximum and minimum values (*n* = 6 larvae). (E,F) Distributions of amplitudes and time constants for fits to step responses calculated using the best (E) and conservative (F) eye position extrapolations. Dots and lines show parameter combinations for larvae for which the best fit was a three-component model (red; best extrapolation: *n* = 5, conservative extrapolation: *n* = 4) or a four-component model (black; best: *n* = 1, conservative: *n* = 2).

Though there are physiological differences between the active and anaesthetized states, step response fits in both cases contained time constants that ranged over several orders of magnitude, including 1 s and 10 s timescales. Quantitatively, the time constants for the active state fits tended to be a few times smaller than for the anaesthetized preparations. However, this may be an artefact of the limited measurement time in the active state; the mean time constants we found when we fit only the first 15 s of post-release eye position in anaesthetized larvae did not differ significantly from the best 3 component fits to active state larvae (p = 0.0832, 0.155, 0.224, 2-sided Wilcoxon rank-sum for each component).

### Oculomotor plant model estimation

We used our step response measurements to compute models of the oculomotor plant that were then used (see “Implications for neural drive”) to estimate the neural drive to the plant in the active state. Because step response measurements reflect both the force applied to the plant, either externally or due to neural drive, and the dynamics of the plant itself, we developed a methodology that enabled us to isolate the plant dynamics. We estimated these dynamics using the standard linear plant model assumption that eye position is the result of the convolution of an applied force with the plant model (i.e. the plant impulse response function). In the case of a known applied force, the plant model can be derived straightforwardly through deconvolution of the eye position response with the applied force. However, in the present case, the applied force is also not known (except for when it equals zero), making this estimation a more challenging “blind deconvolution” problem in which the applied force and plant model must be simultaneously estimated.

To address this problem, we used the fact that, for a multiexponential plant model, the response after release from displacement is a multiexponential decay with time constants equal to those of the plant model (see equation (3)). Previous work has noted that for a displacement long enough to bring all of the mechanical elements in the plant to equilibrium, the amplitude of each exponential component in the step response is proportional to the coefficient of the corresponding plant element (Sklavos et al. 2005). This was used to infer plant model parameters in anaesthetized animals. However, given the time constants we observed in step responses, this equilibrium condition is certainly not met for displacements in the awake animals. Therefore, we implemented a blind deconvolution method to simultaneously infer the parameters of the plant model and the profile of the force applied to the eye (see Methods, Figure 5A). When two responses were recorded from a single larva, the blind deconvolution was performed jointly on both responses so that a single plant model was inferred.

**Figure 5.**
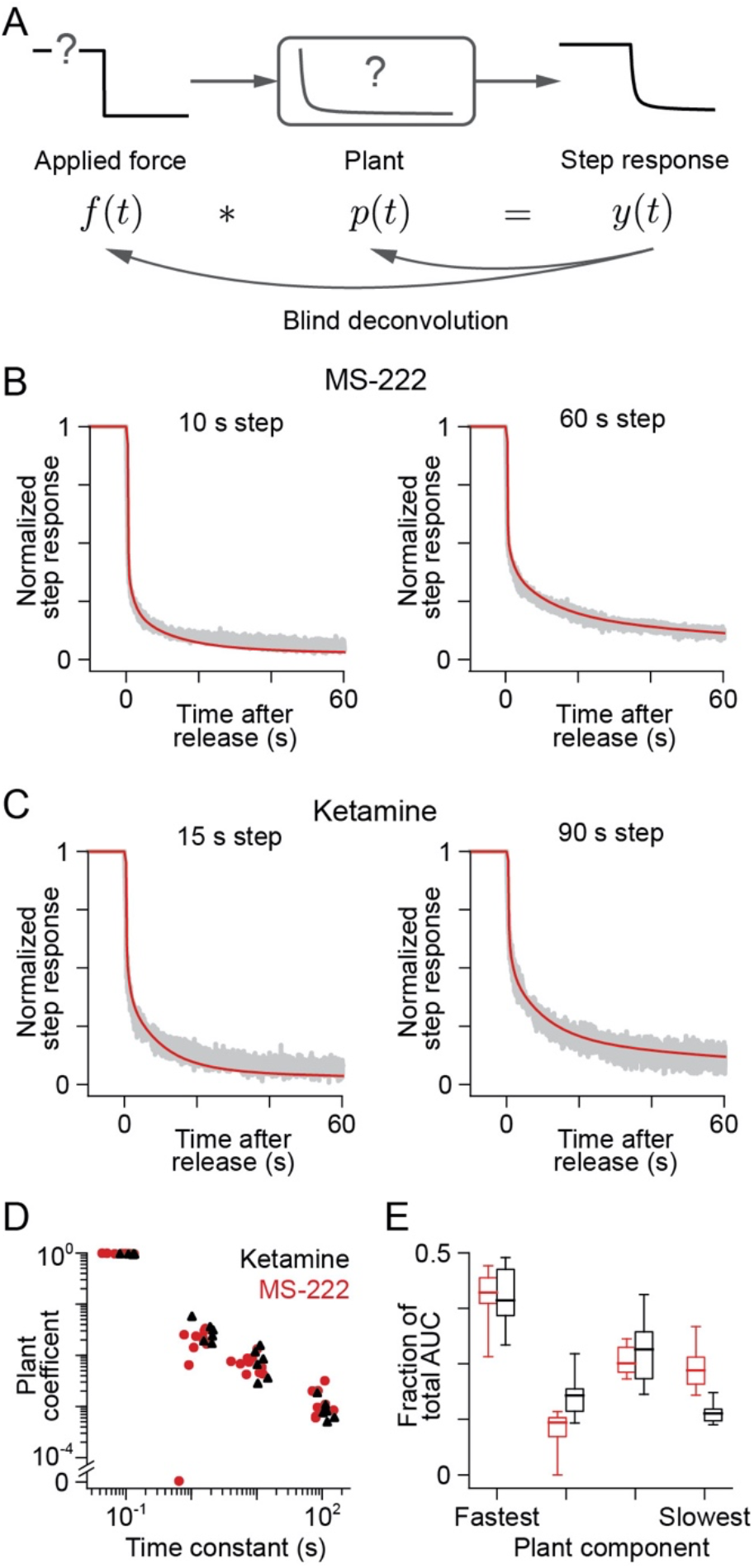
Oculomotor plant model estimates for anaesthetized larvae. (A) Schematic outlining the method used to estimate linear filters that describe plant response properties. We estimated the plant filter and applied force that best fit the measured anaesthetized step responses. (B) Example responses of the eye (grey) following step displacements lasting 10 s (left) or 60 s (right), normalized by the eye position prior to release, in larvae anaesthetized with MS-222, and predicted step response from the best recovered four-component model (red). (C) Same as B, but for larvae anaesthetized with ketamine, for which step displacements lasted 15 s or 90 s. (D) Coefficients and time constants for each component of the best four-component plant models for larvae anaesthetized with MS-222 (red, *n* = 10) or ketamine (black, *n* = 6). (E) Fraction of the total area under the curve (AUC) contributed by each component for the best four-component plant models for larvae anaesthetized with MS-222 (red) or ketamine (black). Boxes span the 25th to 75th percentile range; whiskers show maximum and minimum values (MS-222, *n* = 10; ketamine, *n* = 6).

We first applied our method to the anaesthetized step responses and qualitatively compared our results to previous observations in anaesthetized primates. Using the plant models and applied force profiles inferred by blind deconvolution, we could reconstruct measured eye position reasonably well (Figure 5B,C; mean R^2^ ± sd = 0.870 ± 0.065). We note that the step response reconstructions are less accurate than the direct fits to the step responses performed above (Figure 3A,D), which arises because the plant model was constrained to be identical for the short and long step displacements. This reduced fit accuracy could reflect a small nonlinearity in plant responses or slight nonstationarity between the two recordings. In agreement with previous work in anaesthetized primates (Sklavos et al. 2005), there was an approximately linearly decreasing relationship between the coefficients and corresponding time constants of the inferred plant models on a log-log plot (Figure 5D; parameters of equation (A.6) of Sklavos et al. 2005). Although the long timescale components had relatively small coefficients, due to their long time constants, the integrated area of these components was comparable to that of the faster components (Figure 5E).

We then applied our method to the active state step responses (Figure 6A), obtaining good reconstructions (Figure 6B-D; best extrapolation: mean R^2^ ± sd = 0.936 ± 0.073, conservative extrapolation: mean R^2^ ± sd = 0.960 ± 0.049). As expected from the inverse model framework, the inferred profile of applied force during displacement appeared to be composed of a pulse, step, and some number of exponential slide components (Figure 6C). Like the plant models inferred for anaesthetized larvae, the coefficients of the model components decreased approximately linearly with their corresponding time constants on a log-log plot (Figure 6E), and each component made a comparable contribution to the total area under the curve of the plant model impulse response (Figure 6F).

**Figure 6.**
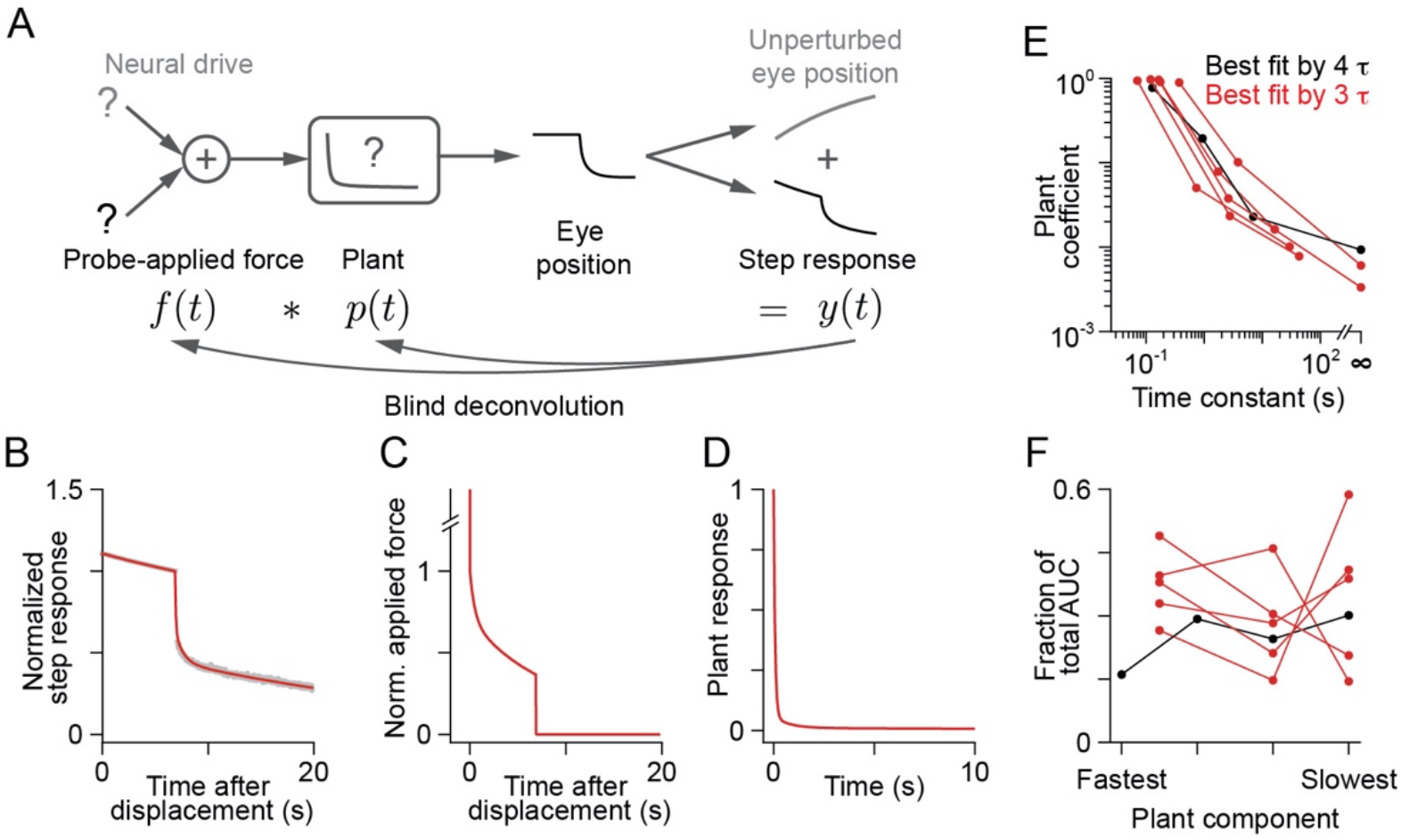
Oculomotor plant models can be recovered from step responses in awake zebrafish larvae. (A) Schematic outlining the method used to estimate linear filters that capture plant response properties. The measured eye position is assumed to derive from the sum of two inputs to the plant: external force applied by the probe, which leads to the active state step response *y*(*t*), and internally generated neural drive, which would lead to the (extrapolated) unperturbed eye position. We estimated the plant filter and applied force that best fit the active state step responses. Active state step responses were estimated from the “best” extrapolation procedure of Figure 2C and normalized to be 1 at the time of release. (B) Example active state step response (grey), and predicted step response from the best recovered three-component plant model (red). (C) Time course of recovered applied force, normalized to be 1 immediately after pulse offset. (D) Time course of the impulse response of the best three-component plant model. (E) Distribution of coefficients and time constants for each component of the best recovered plant models. Dots and lines show parameter combinations for larvae for which a three-component model was best (red, *n* = 5) or for which a four-component model was best (black, *n* = 1). (F) Fraction of the total area under the curve (AUC), calculated over 20 s, contributed by each plant component for the best recovered three- (red, *n* = 5) or four-component (black, *n* = 1) plant models.

### Implications for neural drive

We calculated the neural drive required to stabilize gaze during fixation by deconvolving eye position with the plant impulse response functions inferred from active state step responses (Figure 7A-D). Classical two-element plant models require a neural drive consisting of a brief pulse component, a fast exponential slide component, and a prolonged step component that closely resembles eye position (Optican and Miles 1985; Goldstein and Robinson 1984). For our active state plant models, with component time constants distributed from ~10-100 ms to ~10 seconds, the inferred neural drive instead included multiple exponential slide components and a much smaller step component, so that the amplitude of the neural drive visibly appeared to decay throughout the course of the fixation (Figure 7B, red trace). Solving for the time constants of the neural drive, we found that at least one slide time constant was always > 1 s. For comparison, we calculated the neural drive that would be required given a classical two-element plant with time constants of 20 and 200 ms and coefficients of 0.4 and 0.6, respectively (Robinson et al. 1990). For the classical plant model, the neural drive was essentially constant during fixation (here, we focus on the neural drive > 0.25 s after initiation). Thus, the presence of multiple long timescale components markedly changes the nature of the neural drive required to perfectly stabilize gaze.

**Figure 7.**
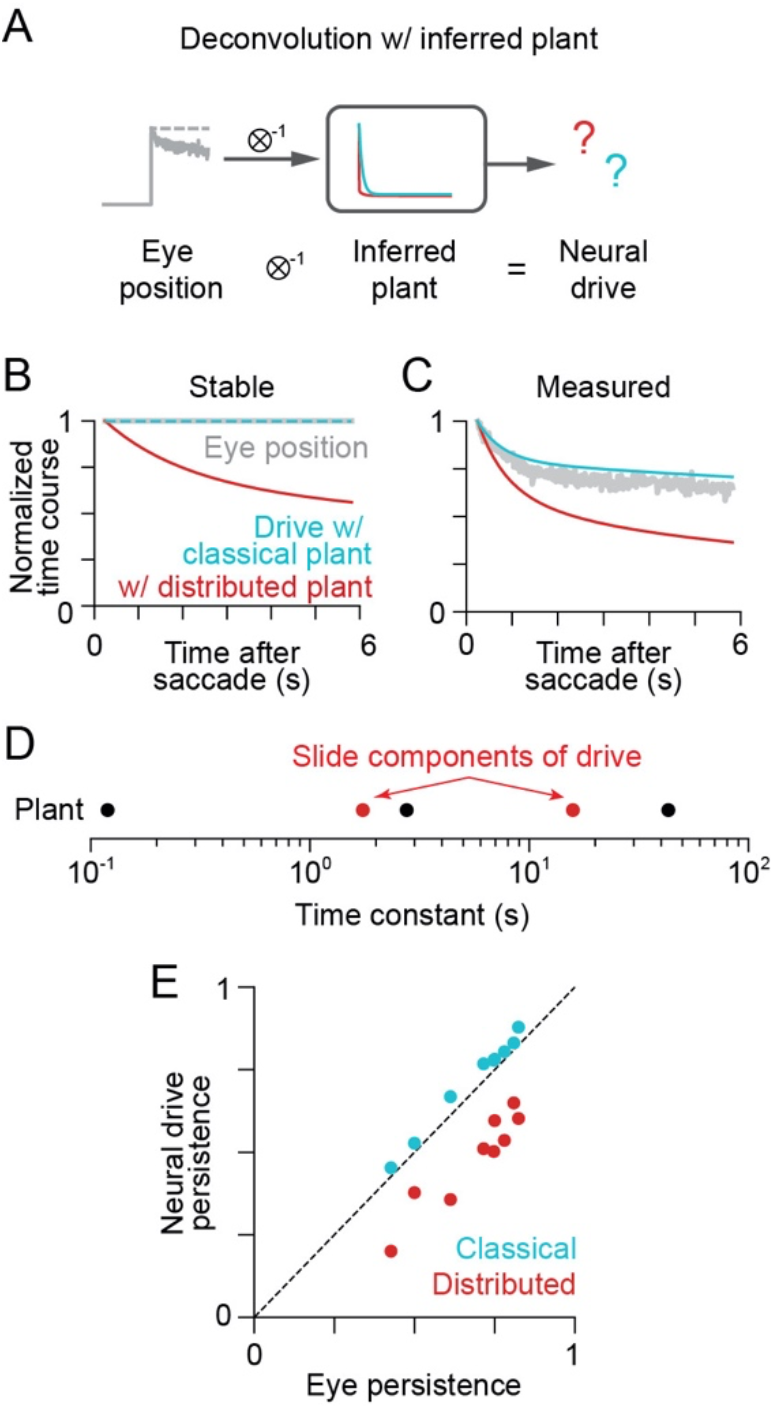
Estimating the neural drive to the oculomotor plant during fixation. (A) Eye position was deconvolved with the filter from a given larva’s plant model to estimate the neural drive needed to generate that eye position. (B) Perfectly stable eye position (grey) and neural drive estimates calculated from the best three-component (red) and classical two-component (cyan) plant model for an example awake larva. Time courses were normalized to be 1 at 250 ms after saccade termination. (C) Same as B, but for a measured fixation. (D) Time constants of the distributed plant (black) and of the slide components of the neural drive (red circles). (E) Persistence of estimated neural drive assuming the best three- or four-component (red) or the classical two-component (cyan) plant model plotted against eye position persistence. Each point represents a fixation recorded from an awake larva. Dotted line indicates a one-to-one ratio of drive persistence to eye persistence.

To understand the nature of the difference in drive for distributed and classical plant models, we analytically derived a formula for the drive as a function of plant model parameters (Mathematical Appendix). This analysis revealed two key differences. First, when long response timescales are present, the drive may no longer be dominated by the step component. Instead, the amplitude of the step component is inversely proportional to the area under the curve of the plant impulse response. As this impulse response becomes longer, the step component’s influence decreases. Second, the presence of multiple plant components that equilibrate over long timescales requires a slowly decaying neural drive. Specifically, we show in the Mathematical Appendix that exactly one slide time constant of the neural drive must fall between each pair of consecutive plant model time constants. This was confirmed by numerical calculation of the slide component time constants for the measured plants (Figure 7D). Thus, the presence of multiple long response timescales necessitates slowly decaying neural drive components.

Actual measurements of larval zebrafish eye position show prolonged but imperfect fixations (Figure 7C, grey trace). Nevertheless, neural drive estimates obtained with distributed plant models, unlike those obtained with classical plant models, again decay more quickly than eye position (Figure 7C, red vs. blue trace). Furthermore, the neural drive contains long timescale slide components identical to those required to maintain perfect fixation (Figure 7D). We show in the Mathematical Appendix that, under the inverse model formulation, the neural drive yielding imperfect fixations has slide components with time constants identical to those of the drive that generates stable fixations, in addition to components resembling recorded eye position that are analogous to the step component for stable fixations.

How much do neural drive estimates for distributed plant models differ from those for classical plant models, given actual eye position measurements? We used “persistence values” to quantify the difference in time course decay expected between neural drive and eye position, assuming either distributed or classical plant models (9 fixations from 6 larvae; Lee et al. 2015). We defined the persistence value for a neural drive or eye position time series as its integral starting from 0.25 s post-saccade, normalized so that a stable time series yields a persistence value of 1. Hence larger persistence values correspond to increasingly persistent time courses. Distributed plant models lead to neural drive persistence that is substantially less than eye persistence, whereas classical plant models imply similar drive and eye persistence (Figure 7E). The median ratios between the persistence values of estimated neural drive and eye position were 0.71 and 1.06 for the distributed and classical plant models, respectively. Thus, distributed plant models predict neural drive will substantially diverge from eye position during fixations.

### The relationship between measured hVPNI activity and eye position

The hVPNI has traditionally been assumed to generate a constant step component, but recent recordings have revealed a diversity of decay timescales in hVPNI firing during fixation (Daie et al. 2015, Miri et al. 2011a). We reanalysed recordings from a previous study (Miri et al. 2011a,b) to evaluate if this heterogeneous activity in the hVPNI is indeed capable of providing the neural drive required to produce observed fixations, given the oculomotor plant models we computed.

We first compared the difference in time course persistence expected between the neural drive and eye position during fixation for individual larvae (Figure 7E) with that seen between measurements of hVPNI firing and simultaneously recorded eye position. Time course persistence was computed for hVPNI neurons whose saccade-triggered average firing rates were estimated from calcium-sensitive cellular fluorescence measured during saccadic eye movement. We modelled saccade-triggered average fluorescence as a multiexponential firing rate function convolved with a calcium impulse response function describing the fluorescence response following an action potential (see Methods, Figure 8A,B; Miri et al. 2011a,b; Daie et al. 2015). Out of an initial dataset of 195 neurons across 6 larvae, we excluded neurons that were not putative integrator neurons (29/195 neurons excluded) such as those exhibiting only bursting activity during saccades, and those for which fluorescence was not well fit by the saccade-triggered average fluorescence model (15/195 neurons excluded), leaving 151 neurons for subsequent analysis. Consistent with previous results, we observed a broad range of persistence values across this population (Figure 8C).

**Figure 8.**
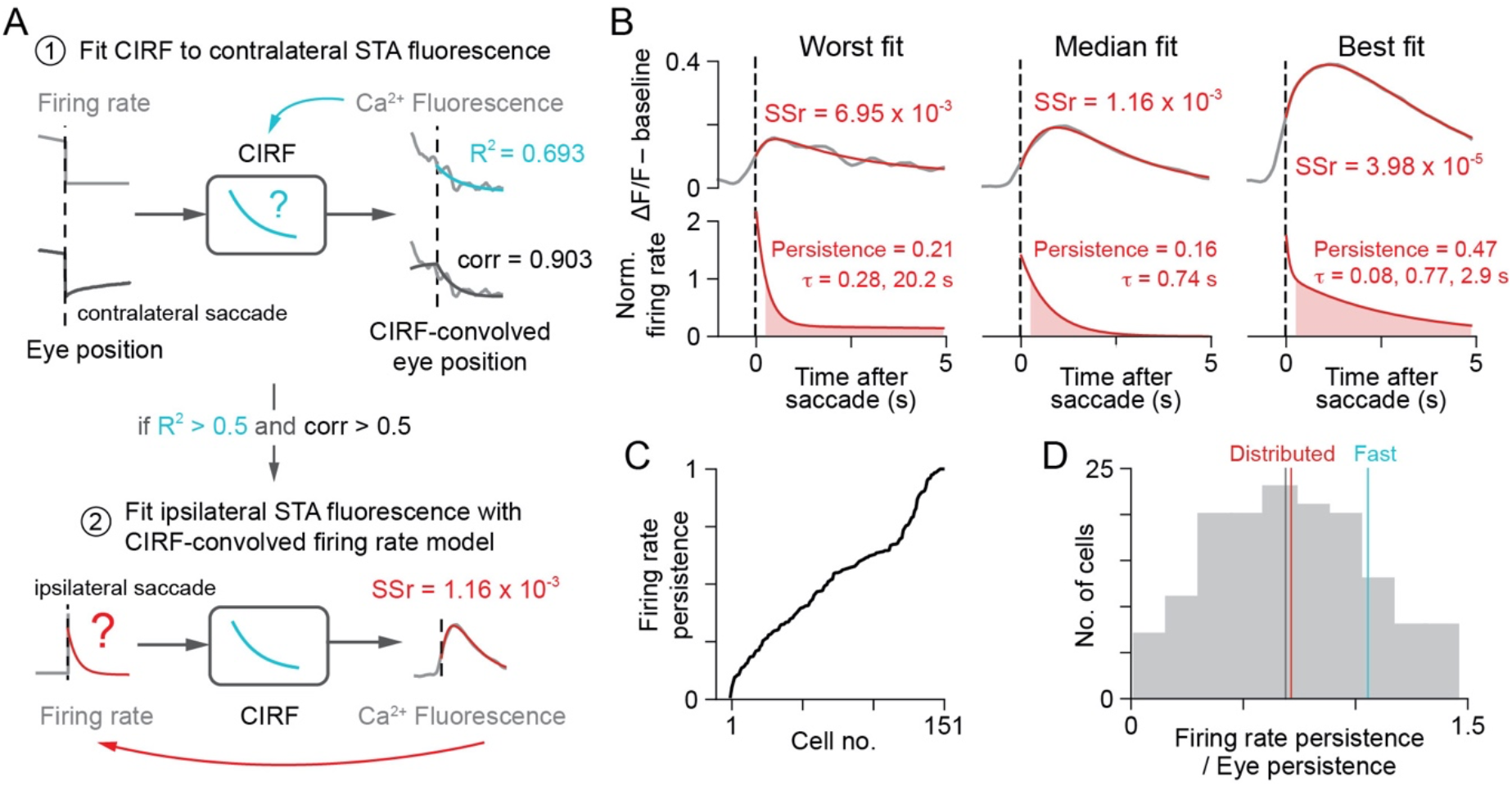
The relationship between the persistence of eye position and of neuronal firing in hVPNI neurons is consistent with the neural drive required for a plant with long timescale responses. (A) Illustration of the method used to recover firing rates of hVPNI neurons. First, the calcium impulse response function (CIRF) was fit to baseline subtracted saccade-triggered average (STA) Ca^2+^-sensitive fluorescence during a contralateral saccade (top). Only cells to which the CIRF fit was relatively good (R^2^ > 0.5), and for which fluorescence was correlated to eye position when convolved with the CIRF (corr > 0.5) were included in further analysis (167/195 cells). For these cells, STA fluorescence during an ipsilateral saccade was fit with a multiexponential model of post-saccadic firing convolved with the CIRF (bottom; Methods, hVPNI firing rate estimation). If the ratio of the sum of squared errors of fits to the sum of squares of ipsilateral STA fluorescence (sum of squares ratio, SSr) was greater than 0.007, the cell was excluded (16/167 cells excluded). (B) Examples of the worst, median and best quality fits to ipsilateral STA fluorescence of included cells (top) and corresponding inferred firing rate functions, normalized to equal 1 at 0.25 s after saccade time (bottom). (C) Distribution of firing rate persistence (sum of red shaded areas in B) across all included neurons. (D) Histogram of the ratio of firing rate persistence to eye position persistence. Grey line indicates the median ratio. Coloured lines indicate the median ratio of neural drive persistence to eye persistence across larvae assuming a distributed (red) or classical (cyan) plant model for each larva.

We then compared persistence values for hVPNI neurons with those of the saccade-triggered average eye position calculated from simultaneously recorded eye position (Figure 8D). We found that firing rate and eye persistence measurements were statistically unlikely to be drawn from the same distribution (p = 0.01106, two-sided Wilcoxon rank-sum). Firing rate persistence was lower than that for the corresponding eye position for 122/151 neurons (81%). The median ratio between the persistence values for firing rate and eye position across all neurons was 0.69, very close to the corresponding value for neural drive estimates (0.71, red line in Figure 8D). Overall, the distribution shown in Figure 8D is more consistent with what would be expected given the distributed plant models than given the classical plant model.

We next addressed whether the firing rates of the population of hVPNI neurons could be used to construct a neural drive that stabilizes gaze given long response timescales in the plant. For this analysis, we generated a three-component summary plant model computed as for the individual larvae but using a simultaneous fit to the active state responses from all six larvae (Figure 9A-C). This summary model had coefficients and time constants similar to the medians of the corresponding distributions from models for individual larvae (Figure 9B). We then deconvolved the saccade-triggered average eye position for each larva with this summary plant model to generate neural drive estimates. Assuming that hVPNI output could effectively be fed forward by motor neurons, we asked whether weighted sums of saccade-triggered average firing rates for recorded hVPNI neurons could well-approximate these neural drive estimates (see Methods, Figure 9D-F). Weights were constrained so that 50% were positive and the rest zero, in agreement with anatomical estimates of the proportion of integrator cells that synapse onto motor neurons (Lee et al. 2015, Vishwanathan et al. 2017). We found that a weighted sum of hVPNI firing could well-approximate the neural drive estimate for all 6 larvae (R^2^ = 1.000). This indicates that hVPNI firing is sufficient to constitute the neural drive needed to stabilize an oculomotor plant characterized by long response timescales.

**Figure 9.**
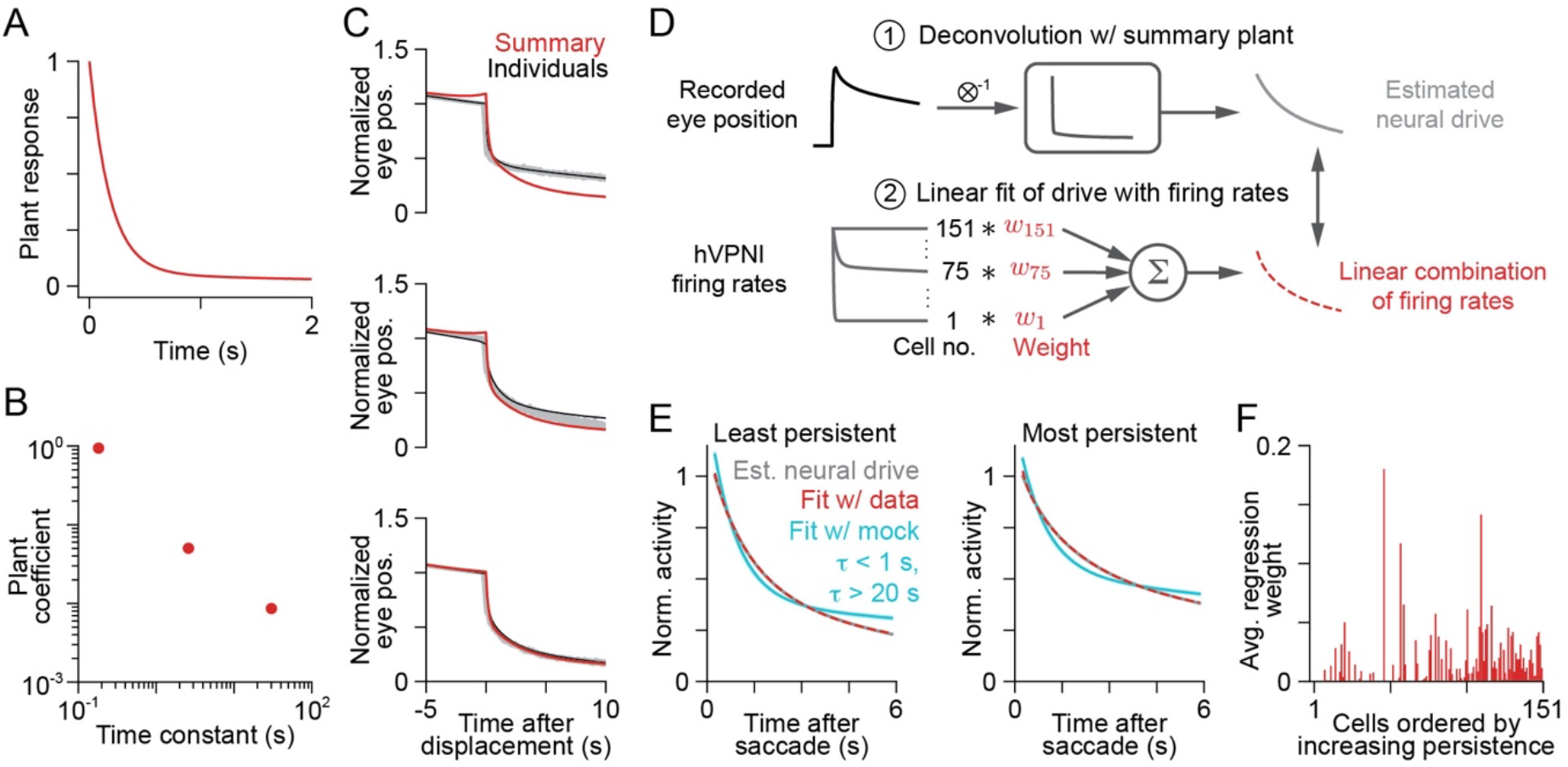
The distribution of hVPNI firing patterns is sufficient to stabilize gaze when the plant has long timescale responses. (A,B) Time course (A) and coefficients and time constants (B) of a single summary plant model fit to the active state step responses from all awake larvae. (C) Predicted step response given the summary plant model (red) compared to the predicted response given the best plant model for each individual larva (black). Examples are the worst (top), median (middle) and best (bottom) of the 9 reconstructed step responses, sorted by mean squared error. (D) Schematic of steps used to estimate the neural drive during fixation as a linear combination of hVPNI firing rates. Saccade-triggered average eye positions during fixation from the 6 larvae from which Ca^2+^-sensitive fluorescence was recorded were deconvolved with the summary plant model to estimate the required neural drive. For each larva we calculated a regularized linear regression of estimated neural drive onto the firing rates of all recorded hVPNI cells (*n* = 151 cells). (E) Grey: estimated neural drive calculated for the fish with the least (left) and most (right) persistent average eye position during fixation using the summary plant model. Red: best linear fit of hVPNI firing rates to the estimated neural drive. Cyan: band containing the best 95% of linear fits to the estimated neural drive from 100 synthetic populations of mock cells whose firing rates were exponential decays with random time constants <1 s or >20 s. (F) Regression weight for each hVPNI cell, averaged across fits to all 6 average eye positions, with cells sorted by increasing firing rate persistence.

That such good fits can be achieved is perhaps not surprising given the range of persistence timescales present in hVPNI activity (Figure 8C). However, we note that intermediate timescales are critical to fit neural drive estimates well. Weighted sums of simulated cell populations containing only a distribution of short (<1 s) and very long (>20 s) timescales could not reconstruct neural drive estimates (Figure 9E, cyan traces; 163- to 978-fold increase in MSE). Thus, the intermediate persistence timescales not present in classical models of the oculomotor neural integrator are a necessary component of this drive.

## DISCUSSION

We report here two significant findings regarding the oculomotor plant and the motor circuits that control it. First, we extend the demonstration of both short (<1 s) and long (>1 s) response timescales in the plant (Quaia et al. 2009; Sklavos et al. 2006; Sklavos et al. 2005) to the active, unanaesthetized state, and to a new vertebrate model organism. Recent reports of long timescale responses in the primate plant have been based on studies using anaesthetized animals. The relevance of these findings to the active state has been argued only indirectly using models (Sklavos et al. 2005). Second, our results establish that, despite such long response timescales, the firing seen among hVPNI neurons (Daie et al. 2015; Miri et al. 2011a) can still be interpreted in terms of an inverse model-based compensation of plant viscoelasticity. While previous work has focused on the encoding of eye position in neuronal populations necessary for gaze stability during fixation (Aksay et al. 2000; Escudero et al. 1992; McFarland and Fuchs 1992; Pastor et al. 1994), a deviation from simply representing eye position appears crucial to the hVPNI’s function. As predicted for an inverse plant model that achieves substantially stable gaze, firing among larval zebrafish hVPNI neurons shows both less persistence on average than eye position itself, and a heterogeneity of persistence timescales.

Previous models of the oculomotor plant that included only short response timescales implied that during fixation, neural drive to the plant would decrease over the first tens or hundreds of milliseconds and thereafter would stably approximate eye position (Goldstein 1984; Optican and Miles 1985; Robinson 1964). This decrease in drive that follows the saccade-inducing burst, attributed to an exponentially decaying slide component, reflects the attenuating force needed to stabilize gaze as the viscoelastic elements equilibrate. The presence of distributed response timescales in the plant that range up to tens of seconds implies a reduced need for the constant, step component of neural drive; instead, distributed timescales of decaying drive extend long into fixations to compensate for the distributed timescales of force dissipation. Indeed, measurements of abducens motor neuron firing during approximately stable fixations in cats show evidence of firing rate decay on timescales greater than 1 s (Davis-Lopez de Carrizosa et al. 2011). Hysteresis observed between abducens motor neuron firing and eye position greater than 2.5 s into fixations (Goldstein and Robinson 1986) is also consistent with the presence of neural drive components on the many seconds timescale, as is the hysteresis seen across timescales between hVPNI neuron firing and eye position (Aksay et al. 2003).

The requirement for neural drive that decays across short and long timescales complicates descriptions of drive to the plant. The notion that the drive is composed of an eye velocity-encoding component, an eye position-encoding component, and an exponentially decaying slide component can be extended to include multiple decaying slide components distributed across a range of timescales (see Mathematical Appendix). But, because sums of exponential components differing in component number, time constants, and amplitudes can well-approximate the same function (Istratov and Vyvenko 1999), discerning precise values for the time constants may not be possible. However, parameters from the multiexponential fits we used to characterize plant responses do meaningfully reflect the breadth of response timescales.

Despite this ambiguity regarding the precise neural drive needed to stabilize gaze, the need for multiple slide timescales does entail a coherent view of the transformation performed by the hVPNI in stabilizing gaze. Each slide timescale can be computed as a “leaky” integral of a brief eye velocity-encoding burst that “leaks” away over time with a particular time constant. The hVPNI can then be viewed as computing a sum of multiple leaky integrals of eye velocity (Figure 1A). Individual hVPNI neurons may reflect distinct combinations of these integrals in their firing, as would the ocular motor neurons they target. Since leaky integration can also be expressed as a convolution with an exponentially decaying filter, the hVPNI can be seen as convolving eye velocity with a multiexponential filter. One practical manifestation of this leaky integration of eye velocity is that hVPNI firing will decay faster than eye position (the pure integral of eye velocity), consistent with our observations of hVPNI firing in the aggregate. While pure integration remains a useful approximation of the transformation performed by the hVPNI, the distributed nature of its integration timescales may have important consequences for the underlying biological mechanisms (Daie et al. 2015; Miri et al. 2011a; Seung 1996; Seung et al. 2000). In previous work, we have identified an array of neural circuit architectures capable of generating leaky integrals on multiple, distributed timescales (Miri et al. 2011a).

This view of the hVPNI as decaying on multiple timescales is conceptually similar to previous work suggesting power law decay of neural firing rates in the oculomotor integrator, since power law decays can be approximated by weighted sums of exponential decay terms. This previous work proposed that the transformation of eye velocity signals by the hVPNI is not pure temporal integration, but fractional-order integration (Anastasio 1994). What is commonly referred to in calculus as integration (integration “of order 1”) can be generalized to integrations of arbitrary real (hence “fractional”) order (Podlubny 1999). Integration of an order between 0 and 1 is equivalent to convolution with a power function filter of the form *t^α^*, where *α* is equal to the integration order minus one. Finding evidence for integration of order between 0 and 1 in oculomotor circuits, Anastasio (1994) posited that the neural drive to the oculomotor plant may constitute a fractional integral of eye velocity. He further proposed that this fractional integral relation might serve to compensate fractional integration of neural drive by the plant. That is, the motor circuitry and the plant itself may each be contributing a fraction of the integration necessary to transform eye velocity signals into a position signal. This view is consistent with suggestions that the oculomotor plant may comprise a broad continuum of response timescales (Sklavos et al. 2005), as relaxation has been observed in the plant on all resolvable timescales. Since a continuous relaxation spectrum, like a power law, can be well-approximated by a sum of exponential terms, our results further bolster this view.

Our results here show that the range of persistence timescales present in hVPNI firing is broad enough to constitute a signal matching estimates of the neural drive needed to dictate eye position during fixation. Although contemporary multiexponential plant models imply that this drive should differ from previous descriptions of hVPNI output emphasizing the encoding of eye position, we show here that linear sums of larval zebrafish hVPNI neuron firing, relayed by motor neurons, could stabilize gaze assuming the plant models we computed. Observations of distributed persistence timescales in the hVPNI of adult goldfish (Major et al. 2004; Miri et al. 2011a) and cat (Davis-Lopez de Carrizosa et al. 2011) suggest that hVPNI output and oculomotor plant dynamics could be similarly reconciled for adult vertebrates.

Despite the sufficiency of hVPNI firing for constituting a drive that can compensate for distributed response timescales in the oculomotor plant, the variation in persistence across neurons may still seem curious. If the plant is well modelled by a single filter, then there will exist a single fixed mapping between eye velocity-encoding commands and the drive needed to produce observed eye position, reflecting a single inverse model of the plant. This single inverse model could be instantiated by the hVPNI such that all hVPNI neurons fired identically, counter to our observations. One possible explanation is that the variation in hVPNI persistence reflects the circuit architecture that gives rise to distributed persistence timescales, or an architecture that additionally enables the context-dependence in the distribution of these timescales that has recently been observed (Daie et al. 2015). Another possibility is that variation in neuronal persistence timescales provides a means for conferring robustness to gaze control. Changes in plant dynamics could be counterbalanced simply by reweighting inputs to motor neurons or the extraocular muscles, effecting a reweighting of different timescales in the neural drive. Lastly, it is also possible that our measurements belie a greater complexity in the plant. It could be that the plant comprises separate dynamic components that are independently compensated by drive to extraocular muscles (Dietrich et al. 2017; Hernandez et al. 2019). This would imply a need for premotor circuitry to instantiate not one but multiple inverse models, and to generate distinct drive components that would not always be in similar proportion. Variation in the firing persistence of hVPNI neurons would then be inevitable.

While gaze stability appears to require something beyond pure temporal integration, the underlying circuitry remains a valuable model for circuit-level short-term memory (Major and Tank 2004). In particular, the multiple timescales of firing persistence seen in the hVPNI appear to be analogous to those seen in cortical short-term memory circuits (Bernacchia et al. 2011; Machens et al. 2010), suggesting that multi-timescale responses may represent a core feature of short-term memory systems throughout the brain. Moreover, models capturing integration on multiple, distributed timescales proposed in previous work on the larval zebrafish hVPNI (Miri et al. 2011a) have been extended to interpret response diversity among oculomotor integrator neurons in monkeys (Joshua et al. 2013). Having established here that that long response timescales are not unique to the primate oculomotor plant, the neural dynamics that compensate for this plant behaviour, and their tuning by the cerebellum, may be further addressed in the larval zebrafish, which is particularly amenable to circuit-level analysis (Ahrens et al. 2012; Arrenberg and Driever 2013; Friedrich et al. 2013; Orger et al. 2008; Miki et al. 2020; Aizenberg and Schuman 2011) and has achieved prominence in the study of visuomotor behaviour (Bianco et al. 2011; Gahtan et al. 2005; Helmbrecht et al. 2018; Okamoto et al. 2008; Portugues and Engert 2009; Sylvester et al. 2017). We note that leaky integration of velocity signals on distributed timescales in the hVPNI still requires the generation of firing that persists much longer than typical membrane and synaptic time constants. Cellular and/or circuit mechanisms must exist that generate this persistence. Thus, elucidating these mechanisms in the larval zebrafish should contribute to our understanding of short-term memory in other circuits.

## DATA AVAILABILITY

Data and code used in this study for data analysis can be found at https://github.com/goldman-lab/oculomotor-response-timescales.

## AUTHOR CONTRIBUTIONS

All experiments were performed in the Tank lab at Princeton University. A.M., E.R.F.A. and D.W.T. conceived of and designed the experiments. A.M. performed the experiments and acquired the data. A.M., B.J.B., and M.S.G. conceived of and performed data analysis and modelling. A.M., B.J.B., E.R.F.A. and M.S.G. drafted the paper. All authors critically revised the paper. All authors approved the final version of the manuscript and agree to be accountable for all aspects of the work in ensuring that questions related to the accuracy or integrity of any part of the work are appropriately investigated and resolved. All persons designated as authors qualify for authorship, and all those who qualify for authorship are listed.

## ACKNOWLEDGEMENTS

We thank Kayvon Daie, Sriram Jayabal, Benjamin Lankow, Alireza Alemi and Jennifer Raymond for helpful conversations.

## MATHEMATICAL APPENDIX

### Analytical calculation of neural drive for a multiexponential plant

Here, we derive analytical expressions for the neural drive expected in the case of a linear multi-exponential plant under the inverse model formulation, extending the work of Sklavos et al. (2005). First, we provide a general formula, then specific formulae for the one- and two-exponential cases, and finally discuss simulation results for three- and four-exponential plants.

Suppose that the measured eye position during normal fixations can be described by sums of *n*_pos_ exponential decays with time constants *τ*_pos,*i*_ = 1/*k*_pos,*i*_ and amplitudes *c*_pos,*i*_,

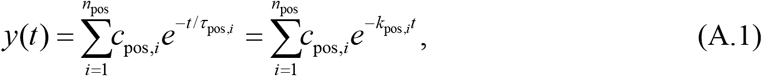

with perfectly stable eye position represented by a single exponential decay with *k*_pos_ = 0. We assume that the eye position is generated by linear filtering of the neural drive by a plant with a multiexponential impulse response function, as in equations (1) and (2). Taking the Laplace transform of equations (1), (2) and (A.1), equation (2) becomes

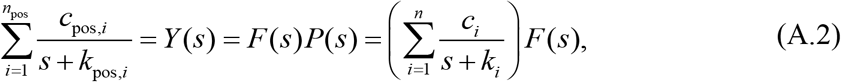

where *k_i_* = 1/*τ_i_* are the inverse time constants, as before, and Laplace transformed variables are represented with capital letters. Since we assume the plant is modelled by Voigt elements in series, we assume the coefficients are positive, *c_i_* > 0. Examining only *P*(*s*), the Laplace transform of *p*(*t*), we can turn the sum of fractions into a single fraction,

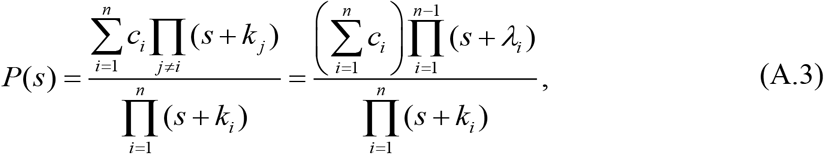

where the numerator is an *n* – 1 degree polynomial with roots −*λ_i_*.

We also write the Laplace transform of the eye position, *Y*(*s*), as a single fraction that is the ratio of two polynomials. Then, resolving the degeneracy in equation (A.2) (see Methods) by assuming that the sum of the coefficients *c_i_* is 1, the Laplace transform of the neural drive is

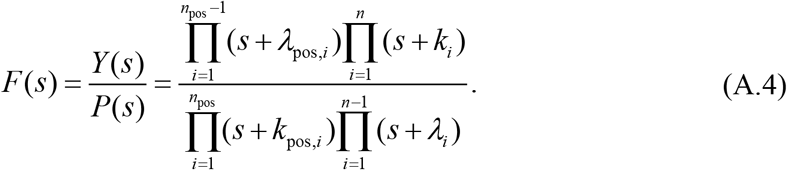

Assume that *λ*_pos,*i*_, *k_i_*, *k*_pos,*i*_ and *λ_i_* are all unique. Then, by partial fraction decomposition, equation (A.4) becomes

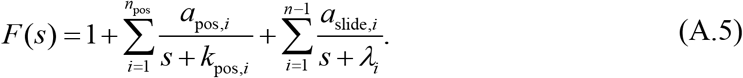

Taking the inverse transform, the neural drive in the time domain is

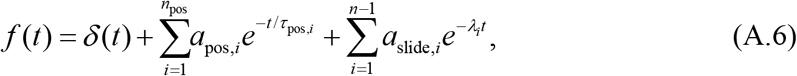

which consists of a pulse, a set of components that decay as eye position, and *n* – 1 exponentially decaying slides with time constants 1/*λ_i_* that may be complicated combinations of the plant coefficients and time constants, but which do not depend on the desired eye position. Note that if the desired eye position had had a component with time constant given by the plant (such that *k*_pos,*i*_ = *k_j_* for some *i* and *j*), then a component with this time constant would not be required in the drive.

From the residue theorem, the amplitude of any eye position component of the drive can be calculated by evaluating (*s* + *k*_pos,*i*_)*F*(*s*) at *s* = –*k*_pos,*i*_,

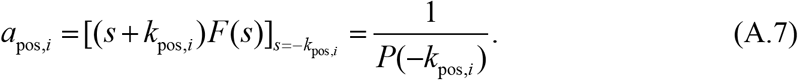

For the special case of perfectly stable eye position, a single component with inverse time constant *k_pos_* = 0, this component is a step with amplitude

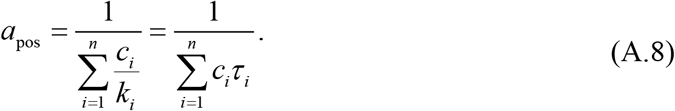

That is, since the area contributed by each plant component *i* is *c_i_ τ_i_*, the amplitude of the step component is equal to the inverse of the total area of the plant filter. To provide additional intuition about the effect of long time constants on the neural drive, we below consider the simple examples of plants with either a single exponential decay or a sum of two exponential decays and for simplicity, eye position that can be described by a single exponential decay.

### A single exponential plant

Consider the case of a single exponential plant model,

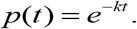

Then, the Laplace transform of the neural drive is

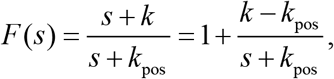

and in the time domain,

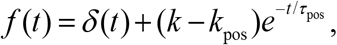

where *k* > *k*_pos_. Thus, in this simplest case, the neural drive consists of a pulse and a decaying component that follows eye position. In the case of stable fixation (*k*_pos_ = 0, *τ*_pos_ → ∞), this component is a step whose amplitude increases in proportion to the decay rate *k* of the plant. To interpret this result, note that convolution with an exponential filter can be thought of conceptually as a leaky integral. To generate stable eye position, an eye velocity command must be supplemented with an eye position command to compensate for the effect of the plant’s leaky filter. As the time constant increases, i.e., as *k* decreases, the plant’s integration becomes less leaky, and the step-like position command becomes less necessary.

### A double exponential plant

Next, consider a plant with two exponential components,

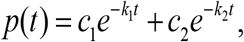

with *c*_1_ + *c*_2_ = 1 and *k*_1_ > *k*_2_, which implies *τ* < *τ*_2_. This has Laplace transform

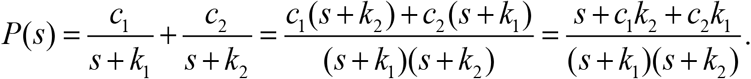

The required neural drive is

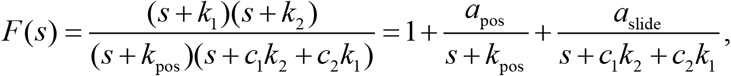

which in the time domain is

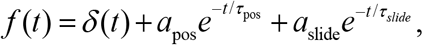

a combination of a pulse, a position-like, and an exponentially decaying slide component, where *a*_pos_ and *a*_slide_ are defined below. The decay rate of the slide is a linear combination of the decay rates of the two plant components,

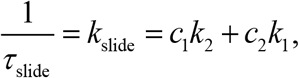

or in terms of time constants,

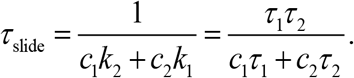

Since *c*_2_ = 1 – *c*_1_, the slide time constant is bounded by the two plant time constants. The slide time constant is longest in the limit where the coefficient of the faster plant component is large (*c*_1_ → 1, *c*_2_ → 0), in which case it approaches *τ*_2_, the time constant of the slower plant component. It is shortest in the limit in which the coefficient of the faster plant component is very small (*c*_1_ → 0, *c*_2_ → 1), in which case it approaches *τ*_2_.

The purpose of the slide can be seen clearly in the case of perfectly stable desired eye position, *k*_pos_ = 0. Applying neural drive with only a pulse and position (step) component to the plant, the eye position due to each plant component *i* is

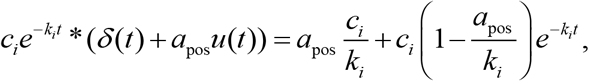

where *u*(*t*) is the Heaviside or unit step function. We can find a step amplitude *a*_pos_ such that the exponential decay term in the above expression is cancelled for each plant component individually, but we will be unable to simultaneously cancel the corresponding exponential decay resulting from the other plant component (a pulse/step mismatch). This is corrected by adding the additional slide component to the drive.

The amplitudes of the position and slide components of the drive are respectively given by

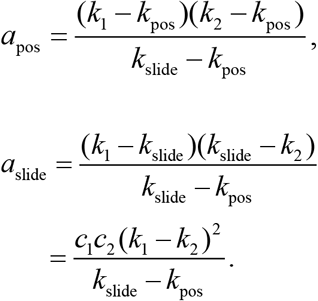

For simplicity, let us consider perfectly stable fixations, *k*_pos_ = 0, so that again the eye position component is a step. Then, the ratio of the amplitude of the slide to the amplitude of the step is given by

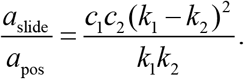

For a fixed choice of time constants, the ratio is proportional to the product *c*_1_ *c*_2_ = *c*_1_, (1 – *c*_1_). If either *c*_1_, or *c*_2_ is sufficiently close to 1, then the amplitude of the slide will be extremely small compared to the step. The ratio is maximized for *c*_1_ = *c*_2_ = 1/2.

In this study, we encountered the situation in which plant components contributed similar areas to the plant, an important parameter setting we will now consider. In the case of two equal area components, with *c*_1_/*k*_1_ = *c*_2_/*k*_2_, the coefficients of the plant components become

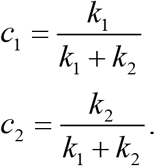

Then, the time constant of the slide is the average of the two plant time constants,

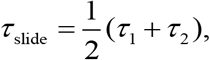

and the amplitudes of the step and the slide are

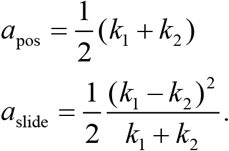

In this case, the contribution of the slide is appreciable only if the inverse time constants of the two plant components are far apart. For the equal areas case, we also can understand easily the effect of a long time constant on the required neural drive. For a fixed choice of time constant *τ*_1_ and coefficient *c*_1_ for the faster component, as the time constant of the slower component becomes longer, the slide time constant increases, the amplitude of the step decreases, and the amplitude of the slide increases.

### General relationship between plant time constants and slide time constants

The observation for the double-exponential plant that the slide time constant of the drive required to stabilize gaze was intermediate between the two plant time constants generalizes to multi-exponential plants. The generalization is that neural drive derived from an inverse model of a multi-exponential plant has slide time constants that are intermediate between neighbouring plant time constants. Thus, long time constants in the plant imply the presence of slowly decaying slides in the drive.

Recall that the time constants of the slide components of the drive *τ*_slide,*i*_ are related to the zeros of the plant –*λ_i_* (the roots of the numerator) as *τ*_slide,*i*_ = 1/*λ_i_*. We will consider a multi-exponential plant with *n* components (equation (A.3)) in which the inverse plant time constants are ordered such that for all *i* ≤ *n*, *k_i_* > *k*_*i*+1_ > 0. We will show that there is exactly one inverse slide time constant *λ_i_* intermediate between each pair of neighbouring inverse plant time constants, i.e., *k_i_* > *λ_i_* > *k*_*i*+1_, as has been previously observed in models of linear viscoelastic materials (Tschoegl 1989).

To show this, first define *P_m_*(*s*) as the first *m* ≤ *n* components of the plant, with time constants ordered as above, i.e.,

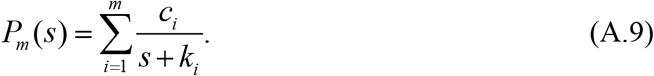

Let *N_m_*(*s*) be the numerator of *P_m_*(*s*) when *P_m_*(*s*) is expressed as a single fraction. Then, we perform proof by induction on *m*. As before, we assume the plant coefficients are positive, *c_i_* > 0.

#### Base case.

For *m* = 2, we have

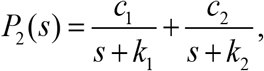

which has numerator

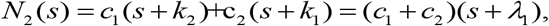

where –*λ*_1_ is the zero of *P*_2_(*s*). We aim to show that *k*_1_ > *λ*_1_ > *k*_2_.

We can write the numerator as the difference of two functions, *N*_2_(*s*) = *f*_2_(*s*) – *g*_2_(*s*), where *f*_2_(*s*) = *c*_1_ (*s* + *k*_2_) and *g*_2_(*s*) = – *c*_2_(*s* + *k*_1_). Then, to find the zero −*λ*_1_, we look for the value of *s* such that *N*_2_(*s*) = 0, i.e., where the lines *f*_2_(*s*) and *g*_2_(*s*) intersect.

Notice that when *s* = −*k*_2_ we have that *f*_2_(−*k*_2_) = 0 and *g*_2_(−*k*_2_) < 0, so that *N*_2_(−*k*_2_) > 0; similarly when *s* = −*k*_1_; we have that *f*_2_(−*k*;) < 0 and *g*_2_(−*k*_1_) = 0, so that *N*_2_(−*k*_1_) < 0. Therefore, by the intermediate value theorem, the intersection point, which is the zero −*λ*_1_, lies in the interval (−*k*_2_, −*k*_1_). Thus, *k*_1_ > *λ*_1_ > *k*_2_, as was to be shown.

#### Inductive step

Now, we assume that the proposition is true for the restricted plant *P*_*m*-1_(*s*), i.e., that each of its zeros 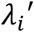 for 1 ≤ *i* ≤ *m* – 2 lies between *k_i_* and *k*_*i*+1_. Then, we have that

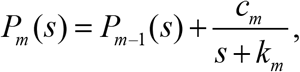

which has numerator

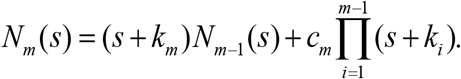

Define *f_m_*(*s*) as the first term of the right hand side, and −*g_m_*(*s*) as the second term, so that *N_m_*(*s*) = *f_m_*(*s*) – *g_m_*(*s*).

First, consider pairs of poles (inverse time constants) of *P*_*m*–1_, i.e., −*k_i_* and −*k*_*i*+1_ for 1 ≤ *i* < *m* – 1. For any of these poles, *g_m_*(−*k_i_*) = *g_m_*(−*k*_*i*+1_) = 0, *k_m_* – *k_i_* < 0 and *k_m_* – *k*_*i*+1_ < 0. Then, we will have that (−*k_i_*) and (−*k*_*i*+1_) will have opposite signs, because by assumption there will be exactly one zero of between −*k_i_* and −*k*_*i*+1_ (so that *P*_*m*–1_ (−*k_i_*) and *N*_*m*–1_(−*k*_*i*+1_) have opposite signs). Therefore, by the intermediate value theorem, *N*_*m*_(*s*) will have at least one zero in the interval (−*k_i_*, −*k*_*i*+1_).

Next, for *s* = −*k*_*m*–1_, we have *g_m_*(−*k*_*m*–1_) = 0. Using that 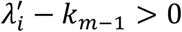 for all *i* < *m* – 1, we also have that *N*_*m*–1_(−*k*_*m*–1_) > 0, and since *k_m_* – *k*_*m*–1_ < 0, we have therefore that *f*_*m*_(−*k*_*m*–1_) < 0. For the added pole *s* = −*k_m_*, we have that *f_m_*(−*k_m_*) = 0, and because *k_i_* – *k_m_* > 0 for all *i* < *m*, we have *g_m_*(−*k_m_*) < 0. Then, *N_m_*(−*k*_*m*–1_) < 0 and *N_m_*(−*k_m_*) > 0, and again by the intermediate value theorem, there is at least one zero in the interval (−*k*_*m*–1_, −*k_m_*).

Since *N_m_*(*s*) is an *m* – 1 degree polynomial, it can have at most *m* – 1 roots. Consequently, since there is at least one root between each of the *m* – 1 pairs of neighbouring poles, *N_m_*(*s*) must have exactly one root (and hence *P_m_*(*s*) exactly one zero) −*λ_i_* in each interval (−*k_i_*, −*k*_*i*+1_) for all 1 ≤ *i* ≤ *m* – 1.

## REFERENCES

Ahrens MB, Li JM, Orger MB, Robson DN, Schier AF, Engert F, and Portugues R. Brain-wide neuronal dynamics during motor adaptation in zebrafish. Nature 485: 471–477, 2012.

Aizenberg M, and Schuman EM. Cerebellar-dependent learning in larval zebrafish. J Neurosci 31(24): 8708–8712, 2011.

Aksay E, Baker R, Seung HS, and Tank DW. Anatomy and discharge properties of pre-motor neurons in the goldfish medulla that have eye-position signals during fixations. J Neurophysiol 84: 1035–1049, 2000.

Aksay E, Major G, Goldman MS, Baker R, Seung HS, and Tank DW. History dependence of rate covariation between neurons during persistent activity in an oculomotor integrator. Cereb Cortex 13: 1173–1184, 2003.

Anastasio TJ. The fractional-order dynamics of brainstem vestibulo-oculomotor neurons. Biol Cybern 72: 69–79, 1994.

Anderson SR, Porrill J, Sklavos S, Gandhi NJ, Sparks DL, and Dean P. Dynamics of primate oculomotor plant revealed by effects of abducens microstimulation. J Neurophysiol 101: 2907–2923, 2009.

Arrenberg AB, and Driever W. Integrating anatomy and function for zebrafish circuit analysis. Front Neural Circuits 7: 74, 2013.

Beck JC, Gilland E, Tank DW, and Baker R. Quantifying the Ontogeny of Optokinetic and Vestibuloocular Behaviors in Zebrafish, Medaka, and Goldfish. J Neurophysiol 92: 3546–3561, 2004.

Becker W, and Klein HM. Accuracy of saccadic eye movements and maintenance of eccentric eye positions in the dark. Vision Res 13: 1021--1034, 1973.

Bernacchia A, Seo H, Lee D, and Wang XJ. A reservoir of time constants for memory traces in cortical neurons. Nat Neurosci 14: 366–372, 2011.

Bianco IH, Kampff AR, and Engert F. Prey capture behavior evoked by simple visual stimuli in larval zebrafish. Front Syst Neurosci 5: 101, 2011.

Blanks RHI, Volkind R, Precht W, and Baker R. Responses of cat prepositus hypoglossi neurons to horizontal angular acceleration. Neurosci 2: 391–403, 1977.

Cohen B, and Komatsuzaki A. Eye movements induced by stimulation of the pontine reticular formation: evidence for integration in oculomotor pathways. Exp Neurol 36: 101–117, 1972.

Daie K, Goldman MS, and Aksay ER. Spatial patterns of persistent neural activity vary with the behavioral context of short-term memory. Neuron 85: 847–860, 2015.

Davis-Lopez de Carrizosa MA, Morado-Diaz CJ, Miller JM, de la Cruz RR, and Pastor AM. Dual encoding of muscle tension and eye position by abducens motoneurons. J Neurosci 31: 2271–2279, 2011.

Dietrich H, Glasauer S, and Straka H. Functional Organization of Vestibulo-Ocular Responses in Abducens Motoneurons. J Neurosci 37: 4032–4045, 2017.

Escudero M, de la Cruz RR, and Delgado-Garcia JM. A physiological study of vestibular and prepositus hypoglossi neurones projecting to the abducens nucleus in the alert cat. J Physiol 458: 539–560, 1992.

Friedrich RW, Genoud C, and Wanner AA. Analyzing the structure and function of neuronal circuits in zebrafish. Front Neural Circuits 7: 71, 2013.

Gahtan E, Tanger P, and Baier H. Visual prey capture in larval zebrafish is controlled by identified reticulospinal neurons downstream of the tectum. J Neurosci 25: 9294–9303, 2005.

Goldman MS, Kaneko CR, Major G, Aksay E, Tank DW, and Seung HS. Linear regression of eye velocity on eye position and head velocity suggests a common oculomotor neural integrator. J Neurophysiol 88: 659–665, 2002.

Goldstein HP, and Robinson DA. Hysteresis and slow drift in abducens unit activity. J Neurophysiol 55: 1044--1056, 1986.

Goldstein HP, and Robinson, D.A. A two-element oculomotor plant model resolves problems inherent in a single-element plant model. Soc Neurosci Abstr 10: 909, 1984.

Green AM, Meng H, and Angelaki DE. A reevaluation of the inverse dynamic model for eye movements. J Neurosci 27: 1346–1355, 2007.

Helmbrecht TO, Dal Maschio M, Donovan JC, Koutsouli S, and Baier H. Topography of a Visuomotor Transformation. Neuron 2018.

Hernandez RG, Calvo PM, Blumer R, de la Cruz RR, and Pastor AM. Functional diversity of motoneurons in the oculomotor system. Proc Natl Acad Sci U S A 116: 3837–3846, 2019.

Istratov AA, and Vyvenko OF. Exponential analysis in physical phenomena. Review of Scientific Instruments 70: 1233–1257, 1999.

Joshua M, Medina JF, and Lisberger SG. Diversity of Neural Responses in the Brainstem during Smooth Pursuit Eye Movements Constrains the Circuit Mechanisms of Neural Integration. J Neurosci 33: 6633–6647, 2013.

Kawato M. Internal models for motor control and trajectory planning. Curr Opin Neurobiol 9: 718–727, 1999.

King WM, Precht W, and Dieringer N. Connections of behaviorally identified cat omnipause neurons. Exp Brain Res 32: 435–438, 1978.

Lee MM, Arrenberg AB, and Aksay ER. A structural and genotypic scaffold underlying temporal integration. J Neurosci 35: 7903–7920, 2015.

Lisberger SG. Internal models of eye movement in the floccular complex of the monkey cerebellum. Neuroscience 162: 763–776, 2009.

Lister JA, Robertson CP, Lepage T, Johnson SL, and Raible DW. nacre encodes a zebrafish microphthalmia-related protein that regulates neural-crest-derived pigment cell fate. Development 126: 3757–3767, 1999.

Machens CK, Romo R, and Brody CD. Functional, but not anatomical, separation of “what” and “when” in prefrontal cortex. J Neurosci 30: 350–360, 2010.

Major G, Baker R, Aksay E, Seung HS, and Tank DW. Plasticity and tuning of the time course of analog persistent firing in a neural integrator. Proc Natl Acad Sci USA 101: 7745–7750, 2004.

Major G, and Tank D. Persistent neural activity: prevalence and mechanisms. Curr Opin Neurobiol 14: 675–684, 2004.

McFarland JL, and Fuchs AF. Discharge patterns in nucleus prepositus hypoglossi and adjacent medial vestibular nucleus during horizontal eye movement in behaving macaques. J Neurophysiol 68: 319–332, 1992.

Miki S, Urase K, Baker R, and Hirata Y. Velocity storage mechanism drives a cerebellar clock for predictive eye velocity control. Sci Rep 10(1): 6944, 2020.

Miri A, Daie K, Arrenberg AB, Baier H, Aksay E, and Tank DW. Spatial gradients and multidimensional dynamics in a neural integrator circuit. Nat Neurosci 14: 1150–1159, 2011a.

Miri A, Daie K, Burdine RD, Aksay E, and Tank DW. Regression-based identification of behavior-encoding neurons during large-scale optical imaging of neural activity at cellular resolution. J Neurophysiol 105: 964–980, 2011b.

Okamoto H, Sato T, and Aizawa H. Transgenic technology for visualization and manipulation of the neural circuits controlling behavior in zebrafish. Dev Growth Differ 50 Suppl 1: S167–175, 2008.

Optican LM, and Miles FA. Visually induced adaptive changes in primate saccadic oculomotor control signals. J Neurophysol 54: 940--958, 1985.

Orger MB, Kampff AR, Severi KE, Bollmann JH, and Engert F. Control of visually guided behavior by distinct populations of spinal projection neurons. Nat Neurosci 11: 327–333, 2008.

Pastor AM, de La Cruz RR, and Baker R. Eye position and eye velocity integrators reside in separate brainstem nuclei. Proc Natl Acad Sci USA 91: 807–811, 1994.

Podlubny I. Fractional differential equations. London: Academic Press, 1999.

Portugues R, and Engert F. The neural basis of visual behaviors in the larval zebrafish. Curr Opin Neurobiol 19: 644–647, 2009.

Quaia C, Ying HS, Nichols AM, and Optican LM. The viscoelastic properties of passive eye muscle in primates. I: static forces and step responses. PLoS One 4: e4850, 2009.

Robinson DA. Integrating with neurons. Ann Rev Neurosci 12: 33--45, 1989.

Robinson DA. The Mechanics of Human Saccadic Eye Movement. J Physiol 174: 245--264, 1964.

Robinson DA. The use of control systems analysis in the neurophysiology of eye movements. Ann Rev Neurosci 4: 463--503, 1981.

Robinson DA, Kapoula Z, and Goldstein HP. Holding the Eye Still After a Saccade. In: Deecke L., Eccles J.C., Mountcastle V.B. (eds) From Neuron to Action. Springer, Berlin, Heidelberg, 1990.

Scudder CA, Kaneko CS, and Fuchs AF. The brainstem burst generator for saccadic eye movements: a modern synthesis. Exp Brain Res 142: 439–462, 2002.

Seung HS. How the brain keeps the eyes still. Proc Natl Acad Sci USA 93: 13339--13344, 1996.

Seung HS, Lee DD, Reis BY, and Tank DW. Stability of the memory of eye position in a recurrent network of conductance-based model neurons. Neuron 26: 259–271, 2000.

Skavenski AA, and Robinson DA. Role of abducens neurons in vestibuloocular reflex. J Neurophysiol 36: 724–738, 1973.

Sklavos S, Dimitrova DM, Goldberg SJ, Porrill J, and Dean P. Long time-constant behavior of the oculomotor plant in barbiturate-d primate. J Neurophysiol 95: 774–782, 2006.

Sklavos S, Porrill J, Kaneko CR, and Dean P. Evidence for wide range of time scales in oculomotor plant dynamics: implications for models of eye-movement control. Vision Res 45: 1525–1542, 2005.

Stahl JS, and Simpson JI. Dynamics of abducens nucleus neurons in the awake rabbit. J Neurophysiol 73: 1383–1395, 1995.

Stahl JS, Thumser ZC, May PJ, Andrade FH, Anderson SR, and Dean P. Mechanics of mouse ocular motor plant quantified by optogenetic techniques. J Neurophysiol 114: 1455–1467, 2015.

Sylvester SJG, Lee MM, Ramirez AD, Lim S, Goldman MS, and Aksay ERF. Population-scale organization of cerebellar granule neuron signaling during a visuomotor behavior. Sci Rep 7: 16240, 2017.

Sylvestre PA, and Cullen KE. Quantitative analysis of abducens neuron discharge dynamics during saccadic and slow eye movements. J Neurophysiol 82: 2612–2632, 1999.

Tschoegl NW. The Phenomenological Theory of Linear Viscoelastic Behavior: An Introduction. Springer, Berlin, Heidelberg, 1989.

Van Opstal AJ, Van Gisbergen JA, and Eggermont JJ. Reconstruction of neural control signals for saccades based on an inverse method. Vision Res 25: 789–801, 1985.

Vishwanathan A, Daie K, Ramirez AD, Lichtman JW, Aksay ERF, and Seung HS. Electron Microscopic Reconstruction of Functionally Identified Cells in a Neural Integrator. Curr Biol 27: 2137–2147 e2133, 2017.

Westerfield M. The Zebrafish Book: A guide for the laboratory use of zebrafish (Danio rerio). Eugene, OR: University of Oregon Press, 2007.

